# Chemogenetic inhibition in the dorsal striatum reveals regional specificity of direct and indirect pathway control of action sequencing

**DOI:** 10.1101/796698

**Authors:** Eric Garr, Andrew R. Delamater

**Affiliations:** Graduate Center, City University of New York; Brooklyn College, City University of New York

## Abstract

Animals engage in intricate action sequences that are constructed during instrumental learning. There is broad consensus that the basal ganglia play a crucial role in the formation and fluid performance of action sequences. To investigate the role of the basal ganglia direct and indirect pathways in action sequencing, we virally expressed Cre-dependent Gi-DREADDs in either the dorsomedial (DMS) or dorsolateral (DLS) striatum during and/or after action sequence learning in D1 and D2 Cre rats. Action sequence performance in D1 Cre rats was slowed down early in training when DREADDs were activated in the DMS, but sped up when activated in the DLS. Acquisition of the reinforced sequence was hindered when DREADDs were activated in the DLS of D2 Cre rats. Outcome devaluation tests conducted after training revealed that the goal-directed control of action sequence rates was immune to chemogenetic inhibition—rats suppressed the rate of sequence performance when rewards were devalued. Sequence initiation latencies were generally sensitive to outcome devaluation, except in the case where DREADD activation was removed in D2 Cre rats that previously experienced DREADD activation in the DMS during training. Sequence completion latencies were generally not sensitive to outcome devaluation, except in the case where D1 Cre rats experienced DREADD activation in the DMS during training and test. Collectively, these results suggest that the indirect pathway originating from the DLS is part of a circuit involved in the effective reinforcement of action sequences, while the direct and indirect pathways originating from the DMS contribute to the goal-directed control of sequence completion and initiation, respectively.

## 1. Introduction

It is well-known that the extent to which an animal’s actions are controlled by the anticipation of future outcomes depends on the functioning of two distinct regions of the striatum: the dorsomedial and dorsolateral striatum (DMS and DLS, respectively; Gremel & Costa, 2013; Yin, Knowlton & Balleine, 2004, 2005; Yin et al., 2005). For example, in the DMS, lesions, NMDA receptor antagonism, and disconnections with prelimbic cortex and basolateral amygdala disrupt instrumental sensitivity to reward devaluation (Balleine, Killcross, & Dickinson, 2003; Gremel & Costa, 2013; Hart, Bradfield, & Balleine, 2018; Yin et al., 2005a, 2005b). Conversely, DLS lesions, rather than interfering with goal-directed control, result in a disruption of habit formation (Gremel & Costa, 2013; Yin, Knowlton, & Balleine, 2004).

Within these two regions of the striatum the balance between two types of neurons—the D1 and D2 receptor-expressing medium spiny neurons (MSNs) of the direct and indirect basal ganglia pathways, respectively—appears to be important in determining the extent to which actions are influenced by their future outcomes. There is evidence to suggest that outcome-sensitive behaviors correlate with activation of D1 MSNs and suppression of D2 MSNs in the DMS, but not the DLS (Furlong et al., 2015; Hong et al., 2019; Li et al., 2015; Shan et al., 2014). In contrast, outcome-insensitive behaviors correlate with suppression of D1 MSNs in the DMS (Furlong et al., 2015) and activation of D1 and D2 MSNs in the DLS (O’Hare et al., 2016).

The studies that have helped to elucidate the behavioral functions of these pathways have been confined to experiments in which animals perform a single action for food rewards. However, theoretical and empirical work suggest that sequences of actions may reveal more nuanced features of goal-directed control (Dezfouli & Balleine, 2012, 2013; Garr & Delamater, 2019). Specifically, Garr and Delamater (2019) showed that the decision to initiate an action sequence following a moderate amount of training was controlled by the anticipation of a future outcome while the execution of the actions within the sequence was not, whereas the reverse was true following an extensive amount of training. That study employed an action sequence task that required rats to press a left lever followed by a right lever for food rewards, and that fixed sequence was continuously reinforced. This type of task differs from the free operant single response tasks that are typically used in studies of striatal correlates of goal-directed control, which require subjects to respond repeatedly on a single manipulandum on a partial reinforcement schedule with few constraints on how sequence of responses are structured (e.g. Corbit et al., 2014; Gremel & Costa, 2013; Lingawi & Balleine, 2012; Yin et al., 2005a, 2005b). The finding reported by Garr and Delamater (2019) that the locus of goal-directed control shifts from sequence initiation to completion over training is not easily captured by any existing model, and, therefore, may lead to additional insights about how the dorsal striatum contributes to the goal-directed control of action sequencing.

In addition to questions about goal-directed control, there is the question of how the direct and indirect pathways contribute to the acquisition and performance of action sequences. There is evidence to suggest that silencing D1 and D2 MSNs in the mouse DLS impedes and enhances sequence performance, respectively (Rothwell et al., 2015). Another study using the same mouse task found that pre-training lesions of the DLS produced deficits in learning a lever-press sequence task (Yin, 2010). Specifically, DLS lesions led to a high rate of perseveration on the right lever and, consequently, a slow rate of learning the left-right sequence, while mice with dorsomedial striatum (DMS) lesions showed a normal rate of learning. However, mice with DLS lesions, but not DMS lesions, also showed prolonged latencies between consecutive lever presses (maximum average of 60 seconds), which raises the possibility that the learning deficit was caused by a performance deficit. If learning the correct sequence of actions requires animals to remember previously performed actions, then animals that move slowly will likely learn at a slow rate because the memory of previously executed actions should decay with time. Indeed, it has been argued that, rather than playing a primary role in instrumental acquisition, the striatum serves to control movement vigor and kinematics (Desmurget & Turner, 2010; Dudman & Krakauer, 2016; Rueda-Orozco & Robbe, 2015; Sales-Carbonell et al., 2018; Thura & Cisek, 2017). On the other hand, the slow latency to complete sequences could have been a product of the slow rate of learning, which could have produced a motivational deficit and, consequently, a slowing of movement.

To more fully investigate the role of the basal ganglia in action sequence learning and performance, we virally expressed Gi-DREADDs (G protein-coupled designer receptors exclusively activated by designer drugs) in either the DMS or DLS and systemically injected clozapine N-oxide (CNO) during and/or after action sequence learning in rats. Gi-DREADDs allow for transient and repeated silencing of neural activity, and can be targeted to specific cell types (see Roth, 2016 for review). In the following experiments, Gi-DREADDs were expressed specifically in striatal neurons associated with either the direct or indirect basal ganglia pathways by using D1 and D2 Cre rats, respectively. Rats were trained to perform a two lever-press sequence identical to a task used in a previous report (Garr & Delamater 2019). This task, which requires rats to perform a specific action sequence to earn food rewards, provides an opportunity to study basal ganglia contributions to sequencing without making inferences about the beginning and end of the sequence, as is sometimes done with free operant, single response tasks (Jin & Costa, 2010; Jin et al., 2014; Matamales et al., 2017; Santos et al., 2015).

## 2. Experiment 1 (inhibiting D1 MSNs in the DMS)

The aim of Experiment 1 was to investigate the effects of suppressing the activity of D1 dopamine receptor-expressing neurons in the DMS during and/or after action sequence learning. Prior to behavioral training, a virus carrying the gene for an inhibitory DREADD or a control virus lacking the gene were virally expressed in the DMS of D1 Cre rats, and rats received injections of either CNO or vehicle before each training session. Validation of DREADD activation by CNO was performed in a separate cohort of rats by combining unilateral DREADD expression, caffeine and CNO injections, and c-Fos immunohistochemistry. Following 20 days of training, rats were subjected to outcome devaluation tests both with and without CNO to measure goal-directed control of action sequences.

### 2.1 Methods

#### 2.1.1 Subjects

Forty-eight naïve Long-Evans rats (22 males and 26 females) were housed in plastic cages (17 x 8.5 x 8 in., l x w x h) in a colony room with a 14-hour light/10-hour dark cycle. Rats were housed in groups of 2 to 4 per cage with wood chip bedding and constant water access. All rats were maintained at 85% of free-feeding body weight for the duration of the experiment by supplemental feedings that occurred immediately following each daily experimental session. Each rat was bred in-house by crossing a D1 Cre transgenic male (source: Rat Resource & Research Center P40OD011062) with a wildtype female (source: Charles River Laboratories). Roughly half of all offspring were confirmed to express Cre in D1 dopamine receptor-expressing neurons (genotyping outsourced to Transnetyx). Only Cre positive rats were used in this experiment.

#### 2.1.2 Apparatus

Eight operant chambers (MED Associates) were used for behavioral training and testing. Each chamber was located within a sound-attenuating box. The interior of the chamber was comprised of two Plexiglas walls, two metal walls, a Plexiglas ceiling, and a grid floor with rods. Attached to one metal wall was a house light. On the opposite metal wall was a food magazine and two retractable levers. The food magazine was connected to two separate pellet dispensers via plastic tubing. The pellets used were TestDiet MLabRodent 45 mg grain pellets and Bio-Serv 45 mg purified pellets. Both pellet types are calorically similar (3.60 and 3.30 kcal/g for Bio-serv and TestDiet, respectively), but have discriminably different tastes. Two lever slots were located to the right and left of the food magazine. Suspended wire cages in a separate room were used for isolating rats during the 1-hour satiation periods, novel pellet pre-exposure, and 20-minute preference tests. During the satiation periods rats were given pellets in ceramic bowls that were stabilized to the cages by hooks attached to springs.

#### 2.1.3 Surgery

Rats were induced with 5% isoflurane and then placed in a stereotaxic frame (Stoelting), where they were maintained on 1-2% isoflurane for the duration of the surgery. Burr holes were drilled in the skull and bilateral infusions were made at the following coordinates relative to bregma: AP: +0.7 mm; ML: +/-2 mm; DV: −5 mm (Paxinos & Watson, 2007). Twenty rats received bilateral infusions of an adeno-associated virus (AAV) carrying the gene for the Gi-DREADD (AAV5-hSyn-DIO-hM4Di-mCherry, titer ≥ 7×10¹² vg/mL, 0.6 µl per side; gift from Bryan Roth; Addgene plasmid #44362; http://n2t.net/addgene:44362; RRID:Addgene_44362) and 20 other rats received bilateral infusions of the control mCherry virus (AAV5-hSyn-DIO-mCherry, 0.6 µl per side; gift from Bryan Roth; Addgene plasmid #50459; http://n2t.net/addgene:50459; RRID:Addgene_50459), counterbalanced with sex and lineage. At the end of each surgery rats were given a subcutaneous injection of buprenorphine (0.05 ml/300 g) and housed in isolation for 5 days before being returned to their original home cages. An additional 8 rats received unilateral infusions of the DREADD AAV (0.8 µl) in the central dorsal striatum for follow-up immunohistochemical verification of DREADD function, and were not part of the main behavioral experiment. The coordinates used, relative to bregma, were: AP, +0.7 mm; ML, +/-2.7 mm; DV, −5 mm. A slightly larger amount of virus was used to cover a wider area in order to express DREADDs across the entire dorsal striatum.

#### 2.1.4 Behavioral training

Rats that received bilateral AAV infusions were trained on an action sequence task beginning a minimum of 3 weeks following surgery. The task was identical to that used in a previous report by Garr and Delamater (2019). Rats were first given magazine training with one pellet type, with half of all rats receiving the TestDiet pellet type and the other half receiving Bio-serv. During this 20-minute session, pellets were delivered according to a 60 second random time schedule, and accompanied by a brief clicker (15 Hz for 0.5 seconds). Rats were then trained to press levers. During the first session of pre-training, the left lever was inserted. A press on the left lever resulted in pellet delivery into the magazine, the retraction of the left lever, and insertion of the right lever. A press on the right lever resulted in pellet delivery into the magazine, the retraction of the right lever, and insertion of the left lever. This cycle continued until 50 pellets were earned or 60 minutes elapsed, whichever occurred first. A second pre-training session was given 24 hours later, in which the conditions were identical to the previous session except that pellets were only delivered following a right lever press.

The main training phase began 24 hours later and continued for 20 daily sessions (Figure 1A). During these sessions, the left and right levers were simultaneously inserted at the beginning of every trial, where they remained inserted until the rat completed a sequence of two lever presses. There were four possible sequences that could be performed: left-left (LL), left-right (LR), right-left (RL), or right-right (RR). If the rat performed an LR sequence, a pellet was delivered and the levers retracted for 1.5 second before being inserted again to start the next trial. If the rat performed any other two-lever sequences pellets were not delivered and the levers retracted for 5 seconds. Thirty minutes prior to each session, rats were given an IP injection of either CNO (source: NIDA; 1 mg/ml/kg, dissolved in 2% DMSO and 98% physiological saline) or vehicle (1 ml/kg, 2% DMSO and 98% saline). CNO solution was made fresh at the beginning of each experimental day. CNO and vehicle injections were balanced with AAV type, such that there were four training groups: DREADD+CNO (n = 10, 5 male and 5 female), DREADD+vehicle (n = 10, 5 male and 5 female), mCherry+CNO (n = 10, 4 male and 6 female), and mCherry+vehicle (n = 10, 4 male and 6 female). Pellet assignment, group assignment, and sex were counterbalanced.

**Figure 1.**
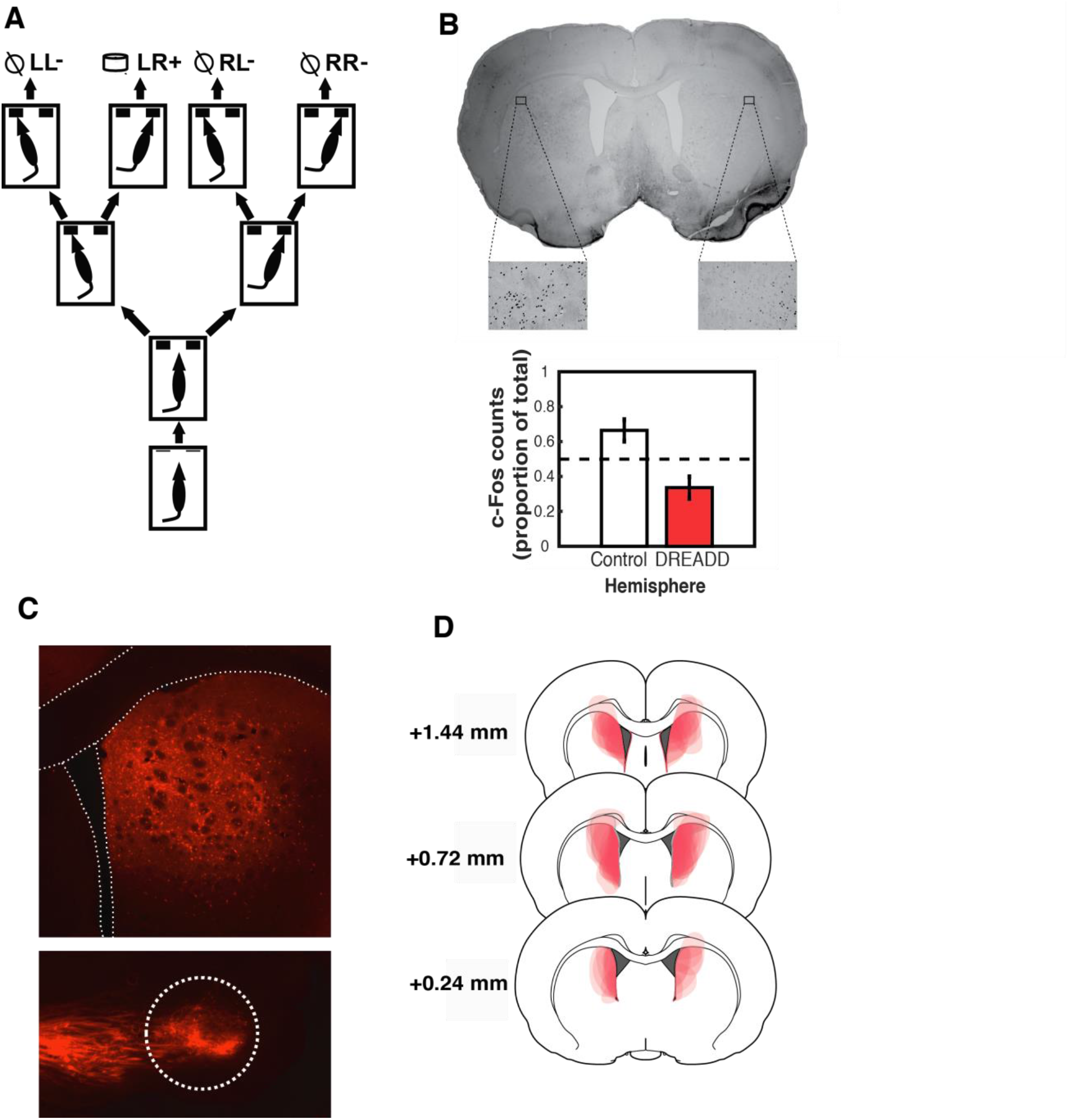
(**A**) Illustration of the action sequence task used in Experiments 1-4. (**B**) Verification of DREADD function. Left: An example coronal section showing c-Fos expression in a D1 Cre rat that received a unilateral Gi-DREADD AAV infusion in the right hemisphere, and was then later given IP injections of CNO followed by caffeine. The insets showing c-Fos positive nuclei are shown at 40x magnification in the DLS for illustration purposes, but note that c-Fos counts were quantified at 10x magnification covering the DLS and central dorsal striatum, where c-Fos was most prominent (see *Methods*). Right: Mean c-Fos counts across control and DREADD hemispheres. (**C**) Top: mCherry expression in a coronal section from a D1 Cre rat that received the DREADD AAV in the DMS. The lateral ventricle and corpus callosum are outlined. Bottom: Sagittal section showing mCherry axon terminal expression in the SNr (**D**) mCherry expression boundaries across all rats given DREADD AAV infusions in the DMS.

#### 2.1.5 Behavioral testing

Two types of tests were conducted following behavioral training. In the first set of tests, animals were given 5-minute extinction tests following injections of CNO and vehicle on different days (order counterbalanced with training group, sex, and pellet assignment). During the tests, the levers operated exactly as they did during training except no pellets were delivered and the clicker was turned off. These tests were designed to separate the effects of DREADD activation on learning versus expression, or both.

The second set of tests were reward devaluation tests. We used the selective satiety procedure in which rats were fed for an hour either on the pellet typed associated with the reinforced sequence or the other pellet type (used as a control for general satiety), and then immediately put through 5-minute non-rewarded tests separated by retraining sessions. All rats received pre-exposure to the novel pellet type the day prior to the start of testing. The pre-exposure procedure consisted of isolating the rats in wire cages until they consumed 20 pellets from a ceramic bowl. Thirty minutes into the satiation sessions, rats were given an injection of either CNO or vehicle. Each rat was tested 8 times: twice after CNO injection and sated on the earned pellet (CNO/devalued), twice after CNO injection and sated on the control pellet (CNO/valued), twice after vehicle injection and sated on the earned pellet (vehicle/devalued), and twice after vehicle injection and sated on the control pellet (vehicle/valued). The order of testing was counterbalanced with AAV type (DREADD vs. control), training injections (CNO vs. vehicle), and sex (male vs. female). One retraining session was run in between each test, during which pellet rewards were reintroduced and injections were given according to the original training conditions.

#### 2.1.6 Histology and immunohistochemistry

Following the end of behavioral testing, rats that were given bilateral AAV infusions and used in the behavioral experiment were perfused transcardially with 0.9% saline followed by 10% formalin. Brains were removed and stored in formalin for 1 hour followed by 30% sucrose in PBS for 72 hours. Coronal sections 40 µm thick were cut using a cryostat, and sections were stored in cryoprotectant at −20 degrees Celsius. A subset of sections from each brain were mounted on microscope slides, coverslipped with Fluoromount (source: Sigma-Aldrich), and examined with a fluorescent microscope (Zeiss).

For rats that were given unilateral AAV infusions and were not part of the main behavioral experiment, after 3 weeks post-surgery they were given an IP injection of CNO (1 mg/kg), followed 30 minutes later by an IP injection of caffeine (100 mg/kg), and then perfused 90 minutes later with 0.9% saline followed by 4% paraformaldehyde. Brains were preserved and sectioned as described above, and sections were then subjected to c-Fos immunohistochemistry. Sections were first rinsed in PBS and then blocked in 3% normal goat serum and 0.25% triton in PBS for one hour. Primary antibody incubation (rabbit anti-c-fos, 1:400) lasted 24 hours. After rinsing in PBS, sections were then incubated in secondary antibody (biotinylated anti-rabbit immunoglobulin, 1:600) for 2 hours, followed by further rinsing in PBS and then incubation in avidin-biotin complex reagent for one hour. Sections were then rinsed in PBS and placed in nickel-intensified diaminobenzidine until sections turned a dark color (no more than 5 minutes). Following a final PBS rinse, sections were mounted on slides and dehydrated in ascending concentrations of ethanol. Slides were coverslipped with Permount and examined with a light microscope (Olympus). c-Fos positive nuclei were counted using a custom macro written in ImageJ (Timothy & Forlano, 2019), and only the central and lateral portions of the dorsal striatum were examined, as these were the regions where c-Fos expression was strongest. Images were taken at 10x magnification.

#### 2.1.7 Statistical analysis

Behavioral measures during training and tests were evaluated using the recommendations of Rodger (1974). This approach treats factorial designs by repartitioning the sum of squares from the standard factorial analysis in order to perform separate one-way ANOVAs (using pooled error terms and Satterthwaite’s (1946) correction for degrees of freedom) to explore the effect of, for example, independent variable A at each level of independent variable B. In addition, the analysis also consists of a main effect test of independent variable B. Significant omnibus F scores are then further examined with a set of ν_1_ mutually orthogonal post-hoc contrasts to determine where differences exist. This approach eliminates the interaction term from the linear model together with the problems associated with interaction tests (see Rodger, 1974). Type I error rate is defined as the proportion of true null contrasts rejected in error, and this is based on Rodger’s table of critical F values (Rodger, 1974). We adopted an α = 0.05 criterion. We also provide a measure of effect size based on Perlman and Rasmussen’s (1975) uniformly minimum variance unbiased estimator of the non-centrality parameter, Δ. When no differences exist in the populations from which samples are drawn, Δ = 0. However, Δ > 0 when true population differences exist. Here we report these estimates whenever significant omnibus F scores were obtained.

### 2.2 Results

#### 2.2.1 Histology and immunohistochemistry

To confirm DREADD function, we used c-Fos immunohistochemistry. It has been shown previously that Gi-DREADD activation by CNO reduces c-Fos counts in the rat dorsal striatum (Ferguson et al., 2011). We induced c-Fos activation by injecting rats with a high dose of caffeine, which has previously been shown to activate c-Fos in the dorsal striatum (Dassesse et al., 1999; Johansson, Lindstrom, & Fredholm, 1994; Svenningsson et al., 1995). Thirty minutes before receiving caffeine injections, rats with unilateral DREADDs received IP injections of CNO. We confirmed that mean c-Fos counts in the dorsal striatum were lower in the DREADD hemisphere compared to the hemisphere with no DREADDs (Figure 1B). A within-subject comparison of normalized c-Fos counts revealed a significant inter-hemispheric difference (*t*(7) = 2.67, *p* < .05). Further one-sample t-tests of normalized counts in each hemisphere against 0.5 chance showed significant differences (*t*’s(7) = 2.63, *p*’s < .05). We conclude that Gi-DREADD activation attenuated cell firing in the dorsal striatum.

For rats that received bilateral DREADD AAV infusions in the DMS and were used in the main behavioral experiment, we observed robust mCherry expression in cell bodies within the DMS (Figure 1D). None of the rats were excluded on the basis of histological analysis. Some brains were also cut in sagittal sections and anterograde expression was examined in SNr, which is a target of D1 MSNs. Fluorescent mCherry expression was confirmed in the SNr (Figure 1C).

#### 2.2.2 Training and expression testing

Before carrying out the main statistical analyses, we first investigated whether there were any sex differences. We only analyzed task acquisition and devaluation test data for this purpose and found no sex differences here or in any of the experiments reported below. All analyses subsequently described are collapsed across sex. We first examined measures of performance during action sequence training. These measures included the total number of sequences performed during each session, and the latency to initiate and complete all sequences during each session. For all training measures, the data were collapsed across the three control groups (DREADD+vehicle, mCherry+CNO, mCherry+vehicle), as there were no statistically significant differences detected among them (between-group ANOVAs performed on every 2-session block: *F*’s < 0.73, *p*’s > .05). Measures were averaged into 2-session blocks because some rats did not provide enough data for a session-by-session analysis. Compared to control rats, the DREADD+CNO group performed significantly fewer sequences during the first 3 blocks of training (Figure 2A; MSE = 2,119.74, *F*’s(1,173) > 5.14, Δ’s > 4.08, *p*’s < .05), were slower to initiate sequences during block 4 (Figure 2A; MSE = 0.59, *F*(1,76) = 6.93, Δ = 5.75, *p* < .05), and were slower to complete sequences during the first 5 blocks (Figure 2A; MSE = 0.38, *F*’s(1,101) > 4.19, Δ’s > 3.11, *p* ‘s < .05). These data indicate that, overall, Gi-DREADD activation in the DMS slowed action sequence performance early in training.

**Figure 2.**
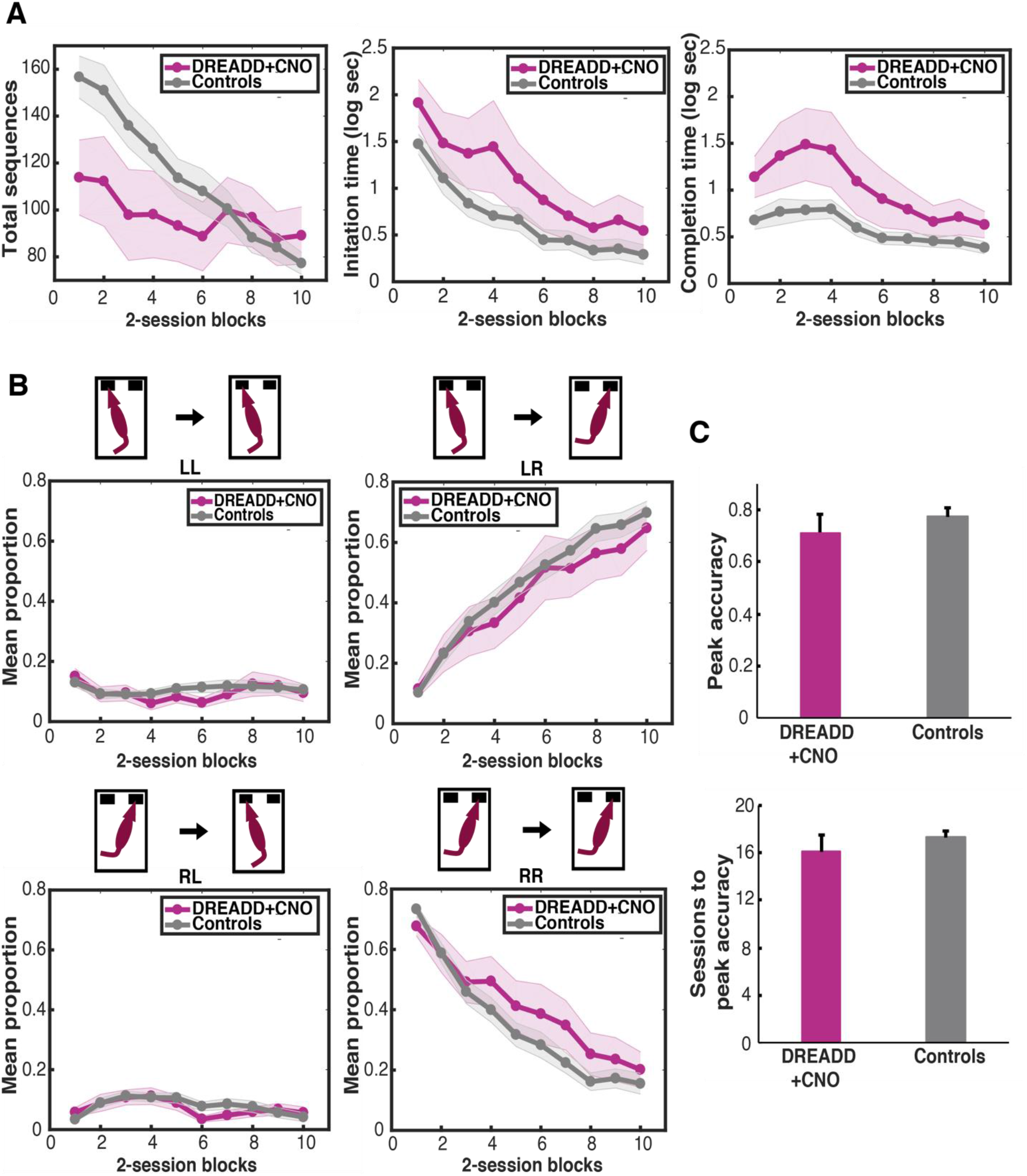
Training data from Experiment 1 (D1 DMS). (**A**) Total sequences (left), initiation times (middle), and completion times (right) across 2-session blocks for rats expressing DREADDs in the DMS and injected with CNO every day prior to training, and controls rats (combined across DREADD+vehicle, mCherry+CNO, and mCherry+vehicle groups). Latency measures are averaged across all sequence types performed within a session. (**B**) Proportions of each sequence type across 2-session blocks. (**C**) Top: Mean peak accuracies for DREADD+CNO and control rats, defined as the maximum proportion of LR sequences achieved in a single session. Bottom: Mean sessions to peak accuracy.

To assess sequence acquisition, the relative proportions of each sequence type were examined (Figure 2B). Once again, data from the three control groups were collapsed, as there were no statistically significant differences detected among the mean LR proportions (between-group ANOVAs performed on every 2-session block: *F*’s(2,63) < 1.06, *p*’s > .05). When comparing controls to DREADD+CNO rats, group means did not differ at any point during training with respect to any sequence type (LL: MSE = 0.01, *F*’s(1,197) < 2.51, *p*’s > .05; LR: MSE = 0.05, *F*’s(1,83) < 0.97, *p*’s > .05; RL: MSE = 0.01, *F*’s(1,141) < 1.89, *p* > .05; RR: MSE = 0.04, F’s(1,197) < 2.82, *p*’s > .05). We also examined peak accuracy, calculated as the maximum proportion of LR sequences achieved in a single session (Figure 2C). There were no group differences when comparing the mean peak accuracies (*t*(37) = 0.90, *p* > .05) or the mean number of sessions to reach peak accuracy (*t*(37) = 0.97, *p* > .05). Thus, Gi-DREADD activation in the DMS did not affect the rate at which an action sequence was learned, although it did slow down overall performance early in training.

During tests of expression conducted after training, there were no detectable within- or between-group differences between CNO and vehicle tests for either total sequences or completion times (*F*’s < 1.15, *p*’s > .05). We conclude that D1 MSNs in the DMS contribute to the speed of sequence performance only early in training.

#### 2.2.3 Devaluation tests

Next, each rat underwent devaluation testing. Sensitivity to devaluation was first assessed by examining the rate of LR sequences under the four testing conditions: CNO/valued, CNO/devalued, vehicle/valued, and vehicle/devalued (Figure 3A). There were no differences between groups during any of the tests (MSE = 35.49; *F*’s(3,121) < 1.43, *p*’s > .05). Collapsing across groups, there was a significant effect of test (MSE = 27.32, *F*(3,105) = 13.64, Δ = 37.14, *p* < .05). Post-hoc contrasts revealed that rats performed fewer sequences during devalued than during valued test sessions after CNO injections (*F*(3,105) = 7.10, *p* < .05) and after vehicle injections (*F*(3,105) = 6.26, *p* < .05). Thus, Gi-DREADD activation did not disrupt goal-directed control of sequences, as measured by the target sequence rate.

**Figure 3.**
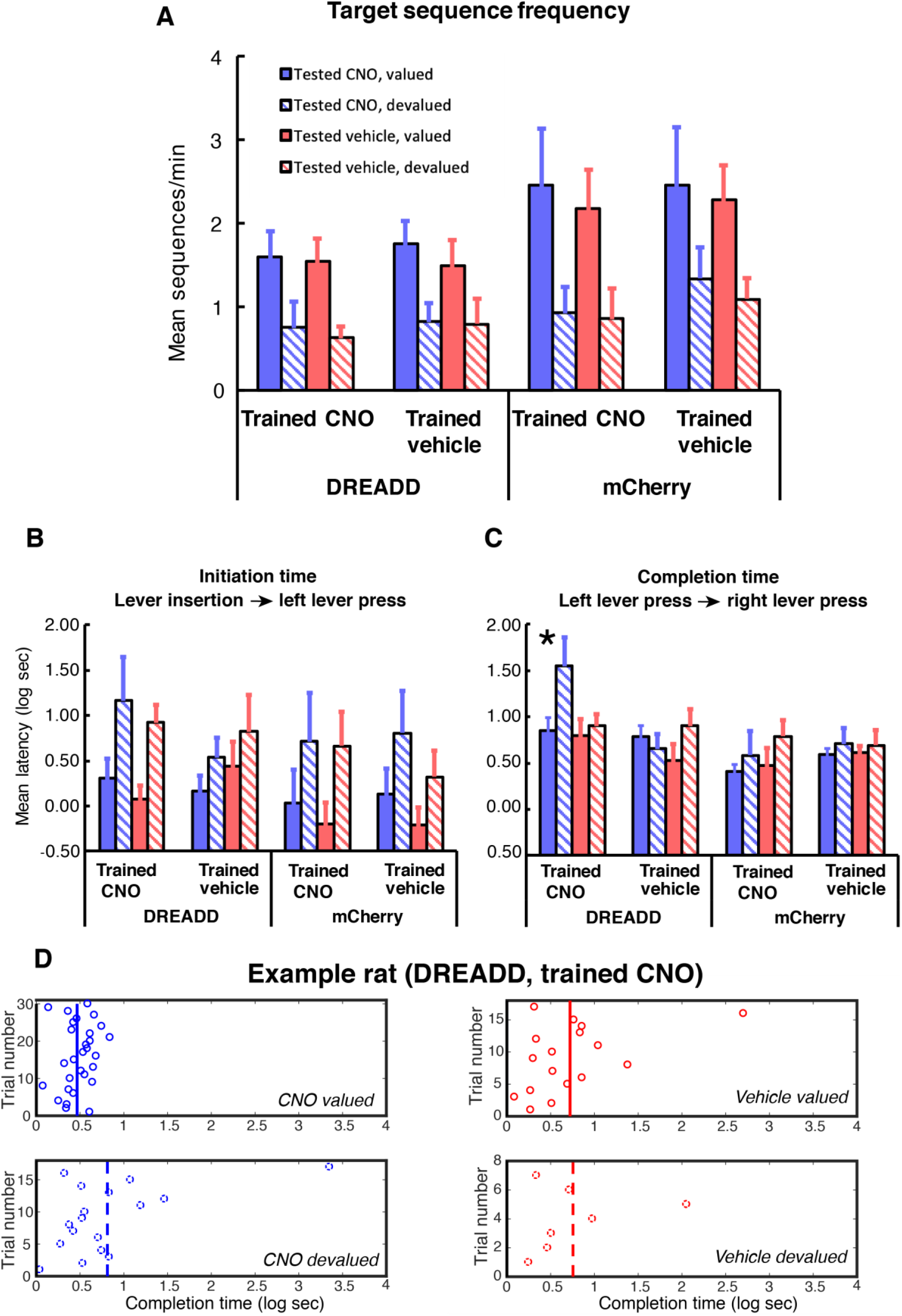
Devaluation test data from Experiment 1 (D1 DMS). (**A**) The rate of LR sequences during devaluation tests. (**B**) Initiation latencies during devaluation tests, defined as the time from lever insertion to the first left lever press. See panel A for legend. (**C**) Completion latencies during devaluation tests, defined as the time from a left lever press to a right lever press during LR trials. See panel A for legend. (**D**) An example rats from the DREADD+CNO training group, showing trial-by-trial LR completion times. Vertical lines represent session means.

We then analyzed latencies to initiate and complete sequences during devaluation tests. For initiation latencies, we analyzed the time from lever insertion to a left lever press (Figure 3B). Since some rats did not generate latency data during one or more of the tests, CNO and vehicle tests were analyzed separately to conserve data. Collapsing across groups, there were significant differences between valued and devalued test sessions, with rats being slower to initiate sequences during devalued test sessions (CNO: MSE = 0.78, *F*(1,33) = 11.43, Δ = 9.74, *p* < .05; Vehicle: MSE = 0.26, *F*(1,31) = 30.10, Δ = 27.16, *p* < .05). There were no differences between groups during any of the tests (*F*’s < 1.27, *p*’s > .05). This analysis further confirms that the chemogenetic manipulation did not affect goal-directed control of sequences, as measured by initiation latency.

For completion latencies, we analyzed the time from a left lever press to a right lever press during LR trials (Figure 3C). Once again, CNO and vehicle tests were analyzed separately. A between-group ANOVA revealed a group difference during the CNO/devalued tests (MSE = 0.29, *F*(3,64) = 6.37, Δ = 15.51, *p* < .05), and post-hoc contrasts revealed a longer latency for the DREADD+CNO group compared to all other groups (*F*(3,64) = 6.35, *p* < .05), which themselves did not differ. Groups were equally quick to complete LR sequences during CNO/valued test sessions (MSE = 0.29, *F*(3,64) = 1.14, *p* > .05). No between-group differences were detected during vehicle test sessions. When collapsing across groups, there were no significant overall differences between valued and devalued test sessions (CNO: MSE = 0.30, *F*(1,32) = 2.53, *p* > .05; Vehicle: MSE = 0.20, *F*(1,29) = 3.61, *p* > .05). These analyses show that chemogenetic inhibition during training and test slowed completion latencies when rewards were devalued, but otherwise completion latencies were insensitive to devaluation. A representative set of individual rat data is shown in Figure 3D.

Consumption data from the satiation periods showed that rats consumed the same amount of pellets across the four different test conditions (MSE = 24.31, *F*’s(3,105) < 1.80, *p*’s > .05). There were no between-group differences (MSE = 45.29, *F*(3,35) = 0.32, *p* > .05). To assess whether the satiation period induced selective satiety, preference tests were conducted following the extinction tests. A preference score was calculated as the percent preference for the pellet type that the rats were not exposed to during the satiation period, calculated separately for CNO and vehicle tests. Within-group ANOVAs on CNO and vehicle preference scores revealed no significant differences (MSE = 0.01, *F*’s(1,34) < 0.81, *p*’s > .05). There were also no between-group differences (MSE = 0.04, *F*(3,34) = 0.62, *p* > .05). Collapsing across CNO and vehicle tests, the mean preference scores for the DREADD+CNO, DREADD+vehicle, mCherry+CNO, and mCherry+vehicle groups were 95%, 87%, 94%, and 94%, respectively. These analyses show that rats were not differentially sated during the four different tests, and that the satiety treatment was selective.

### 2.3 Discussion

We sought to determine whether D1 MSNs in the DMS are necessary for action sequence learning and performance as well as its goal-directed control. There were four main findings. First, Gi-DREADD activation slowed sequence performance early in training, as measured by the number total sequences performed, and the latency to initiate and complete sequences. Second, Gi-DREADD activation did not affect the rate at which a reinforced sequence was acquired. Third, Gi-DREADD activation did not alter goal-directed control of the previously reinforced sequence, as measured by the sequence rate and the latency to initiate sequences during selective satiation tests. Fourth, and finally, Gi-DREADD activation slowed completion latencies during outcome devaluation, but only for rats that received CNO during training and test. Interpretation of these results are deferred to the *General Discussion*.

## 3. Experiment 2 (inhibiting D1 MSNs in the DLS)

### 3.1 Methods

#### 3.1.1 Subjects

Forty-two naïve Long-Evans rats (24 males and 18 females) were housed in identical conditions as rats in Experiment 1. Each rat was bred by crossing a D1 Cre transgenic male (source: Rat Resource & Research Center P40OD011062) with a wildtype female (source: Charles River Laboratories). Roughly half of all offspring were confirmed to express Cre in D1 dopamine receptor-expressing neurons (genotyping outsourced to Transnetyx). Only Cre positive rats were used in this experiment.

#### 3.1.2 Apparatus

The apparatus was identical to that used in Experiment 1.

#### 3.1.3 Surgery

Rats underwent stereotaxic surgery in which the same AAV’s as in Experiment 1 were bilaterally infused, but at the following coordinates (relative to bregma): AP, +0.7 mm; ML, +/-3.6 mm; DV, −5 mm (Paxinos & Watson, 2007). The method of surgery was the same as in Experiment 1. Twenty-two rats received bilateral infusions of the AAV carrying the gene for the Gi-DREADD and 20 other rats received bilateral infusions of the control mCherry virus, counterbalanced with sex and lineage.

#### 3.1.4 Behavioral training

Rats were trained on the same action sequence task as that used in Experiment 1 for 20 daily sessions, beginning a minimum of 3 weeks following surgery. There were four groups: DREADD+CNO (*n* = 11, 6 male and 5 female), DREADD+vehicle (*n* = 11, 6 male and 5 female), mCherry+CNO (*n* = 10, 6 male and 4 female), and mCherry+vehicle (*n* = 10, 6 male and 4 female). Pellet assignment, group assignment, and sex were counterbalanced.

#### 3.1.5 Behavioral testing

Expression tests and devaluation tests proceeded exactly as in Experiment 1.

#### 3.1.6 Histology

Rats were perfused and brains were sectioned and imaged exactly as in Experiment 1.

### 3.2 Results

#### 3.2.1 Histology

We observed robust mCherry expression in cell bodies within the DLS. Two rats (one in the DREADD+CNO group and one in the DREADD+vehicle group) were excluded from all analyses because fluorescence extended into the DMS. The boundaries of fluorescent expression for all other rats are presented in figure 4A.

**Figure 4.**
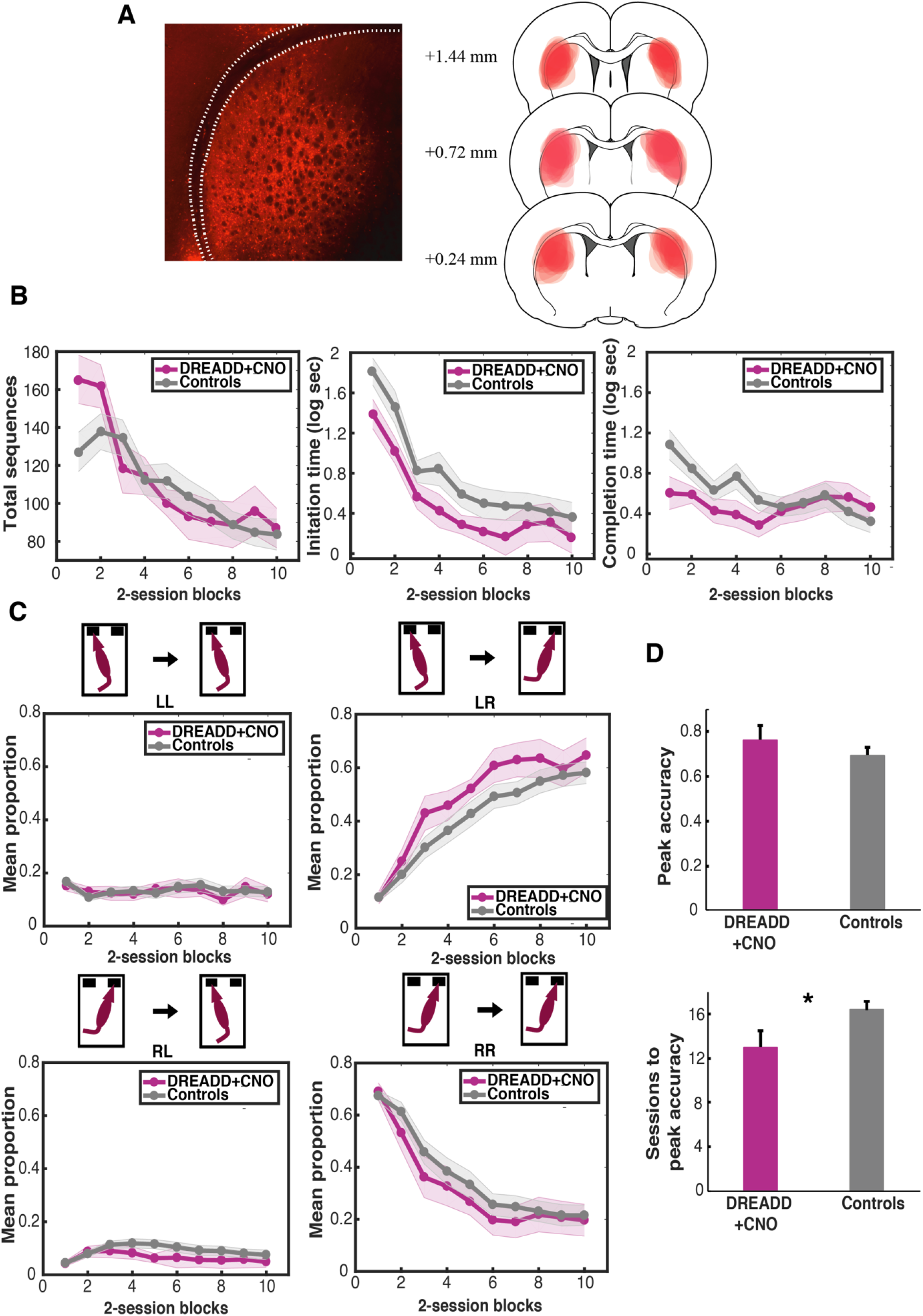
Training data from Experiment 2 (D1 DLS). (**A**) Left: Example coronal section showing mCherry expression in the DLS. The corpus callosum is outlined. Right: mCherry expression boundaries across all rats given DREADD AAV infusions in the DLS. (**B**) Total sequences (left), initiation times (middle), and completion times (right) across 2-session blocks for rats expressing DREADDs in the DLS and injected with CNO every day prior to training, and controls rats (combined across DREADD+vehicle, mCherry+CNO, and mCherry+vehicle groups). Latency measures are averaged across all sequence types performed within a session. (**C**) Proportions of each sequence type across 2-session blocks. (**D**) Top: Mean peak accuracies for DREADD+CNO and control rats, defined as the maximum proportion of LR sequences achieved in a single session. Bottom: Mean sessions to peak accuracy.

#### 3.2.2 Training and expression testing

We once again examined measures of performance during training by analyzing the total number of sequences performed during each session, and the latency to initiation and complete all sequences during each session. Compared to control rats, the DREADD+CNO group performed significantly more sequences during the first block of training (Figure 4B; MSE = 2,043.73, *F*(1,142) = 5.35, Δ = 4.27, *p* < .05), and were also faster to complete sequences during the first block (Figure 4B; MSE = 0.35, *F*(1,111) = 4.95, Δ = 3.86, *p* < .05). There were no between-group differences detected with respect to initiation times (MSE = 0.53, *F*’s(1,89) < 2.63, *p*’s > .05). These data indicate that, overall, Gi-DREADD activation in the DLS facilitated action sequence performance early in training.

To assess sequence acquisition, the relative proportions of each sequence type were examined (Figure 4C). Data from the three control groups were collapsed, as there were no statistically significant differences detected among the mean LR proportions (between-group ANOVAs performed on every 2-session block: *F*’s(2,57) < 0.79, *p*’s > .05). It appears as though the DREADD+CNO group may have acquired the LR sequence more rapidly and gave up repetitive RR sequences more rapidly than the control groups. However, when comparing controls to DREADD+CNO rats at each training block, no significant differences were detected with respect to any sequence type at any training block (LL: MSE = 0.01, *F*’s(1,108) < 0.30, *p*’s > .05; LR: MSE = 0.04, *F*’s(1,87) < 2.53, *p*’s > .05; RL: MSE = 0.01, *F*’s(1,142) < 3.03, *p*’s > .05; RR: MSE = 0.05, F’s(1,102) < 1.45, *p*’s > .05). We also examined peak accuracy (Figure 4D), which revealed no group difference (*t*(37) = 0.94, *p* > .05). However, the number of sessions to reach peak accuracy revealed a group difference consistent with the DREADD+CNO group requiring fewer sessions to reach peak accuracy (*t*(37) = 2.13, *p* < .05). These analyses provide partial support for the suggestion that Gi-DREADD activation in the DLS facilitated the learning of a reinforced action sequence, although this effect was small and did not show up reliably on all measures of learning. It is possible that the modest speeding up of sequence learning was caused by the speeding up of sequence performance induced by Gi-DREADD activation, although a correlation between total sequences performed during the first training block and number of sessions to peak accuracy did not yield significant correlation coefficients (*r*’s = −0.36 and −0.04 for DREADD+CNO and controls, respectively; *p*’s > .05).

During tests of expression conducted after training, there were no detectable within- or between-group differences between CNO and vehicle tests for either total sequences or completion times (*F*’s < 1.88, *p*’s > .05). We conclude that D1 MSNs in the DLS contribute to the speed of sequence performance only early in training.

#### 3.2.3 Devaluation tests

Sensitivity to devaluation was first assessed by examining the target sequence rate under the four testing conditions: CNO/valued, CNO/devalued, vehicle/valued, and vehicle/devalued (Figure 5A). Overall, each group displayed more LR sequences during valued than devalued test sessions with both CNO and Vehicle, but the groups did not differ in this regard. Again, in order to avoid losing data (from missing cells) the CNO and vehicle data were analyzed separately. There were no differences between groups during any of the tests (MSE = 38.92; *F*’s(3,121) < 0.42, *p*’s > .05). However, collapsing across groups, there was a significant effect of test (MSE = 29.80, *F*(3,105) = 12.24, Δ = 33.02, *p* < .05), and post-hoc contrasts revealed that rats performed fewer sequences during devalued than during valued test sessions after CNO injections (*F*(3,105) = 5.40, *p* < .05) and also after vehicle injections (*F*(3,105) = 6.85, *p* < .05). Thus, Gi-DREADD activation in the DLS did not disrupt goal-directed control of sequences, as measured by the target sequence rate.

**Figure 5.**
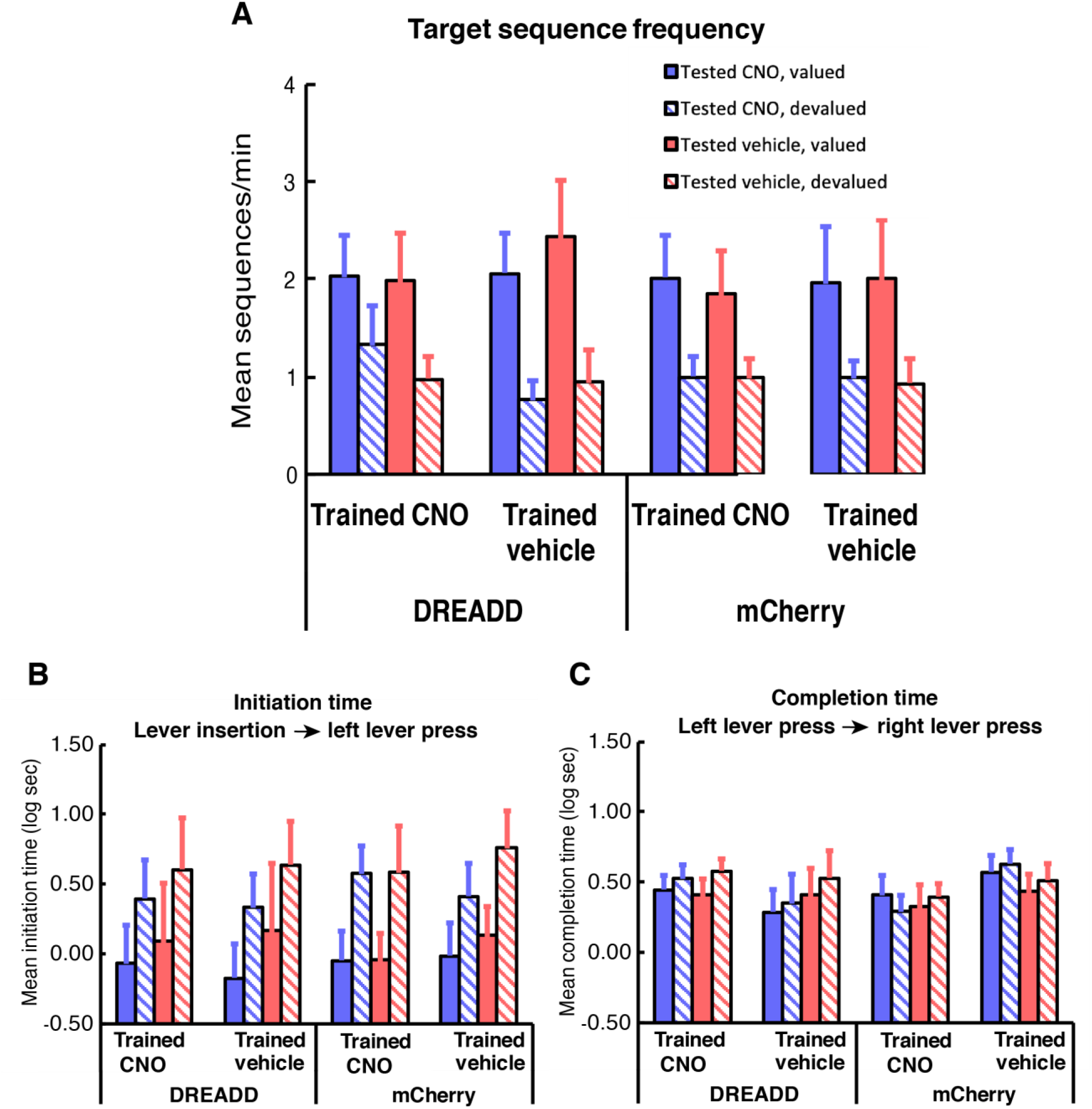
Devaluation test data from Experiment 2 (D1 DLS). (**A**) The rate of LR sequences during devaluation tests. (**B**) Initiation latencies during devaluation tests, defined as the time from lever insertion to the first left lever press. See panel A for legend. (**C**) Completion latencies during devaluation tests, defined as the time from a left lever press to a right lever press during LR trials. See panel A for legend.

We then analyzed latencies to initiate and complete sequences during devaluation tests as in Experiment 1 (Figure 5B). CNO and vehicle tests were once again analyzed separately because some rats did not perform an LR sequence in one of the test sessions. There were no differences between groups during any of the tests (*F*’s < 0.18, *p*’s > .05). Collapsing across groups, there were significant differences between valued and devalued test sessions, with rats being slower to initiate sequences during devalued test sessions (CNO: MSE = 0.28, *F*(1,31) = 15.82, Δ = 13.80, *p* < .05; Vehicle: MSE = 0.41, *F*(1,35) = 14.52, Δ = 12.69, *p* < .05). This analysis further confirms that the chemogenetic manipulation did not affect goal-directed control of sequences, as measured by initiation latency.

For completion latencies (Figure 5C), there were no between-group differences detected during any of the tests (CNO: MSE = 0.14, *F*’s(3,38) < 1.52, *p*’s > .05; Vehicle: MSE = 0.19, *F*’s(3,56) < 0.30, *p*’s > .05) and collapsing across groups, there were no significant differences between valued and devalued test sessions (CNO: MSE = 0.03, *F*(1,30) = 0.17, *p* > .05; Vehicle: MSE = 0.10, *F*(1,34) = 2.27, *p* > .05). These analyses show that chemogenetic inhibition spared the insensitivity of sequence completion times to outcome devaluation.

Consumption data from the satiation periods showed that rats from the DREADD+CNO, DREADD+vehicle, and mCherry +vehicle groups consumed the same amount of pellets across the four different test days (MSE = 15.26, *F*’s(3,105) < 1.15, *p*’s > .05). The mCherry+CNO group consumed different amounts of pellets across the four tests (F(3,105) = 2.44, Δ = 4.18, *p* < .05), and post-hoc contrasts revealed greater consumption during the devalued versus valued tests days collapsing over injection type (*F*(3,105) = 2.26, *p* < .05). There were no between-group differences in overall intakes (MSE = 55.88, *F*(3,35) = 0.58, *p* > .05). Within-group ANOVAs on CNO and vehicle preference scores revealed no significant differences for DREADD+CNO, DREADD+vehicle, or mCherry+CNO groups (MSE = 0.01, *F*’s(1,35) < 2.43, *p*’s > .05). The mCherry+vehicle group showed a significantly greater preference score during CNO compared to vehicle tests (91% vs. 83%, *F*(1,35) = 5.83, Δ = 4.50, *p* < .05). Collapsing across CNO and vehicle tests, the mean preference scores for the DREADD+CNO, DREADD+vehicle, mCherry+CNO, and mCherry+vehicle groups were 96%, 96%, 92%, and 92%, respectively. There were significant differences among the mean scores (MSE = 0.02, *F*(1,35) = 2.05, *p* < .05), with the two DREADD groups showing a greater preference for the non-sated pellet type.

### 3.3 Discussion

We sought to determine whether D1 MSNs in the DLS are necessary for action sequence learning and performance. There were four main findings. First, Gi-DREADD activation sped up sequence performance early in training, as measured by the number of total sequences performed, and the latency to complete sequences. Second, Gi-DREADD activation facilitated the rate at which a reinforced sequence was acquired by some, but not all measures, although this was likely a consequence of rats performing more sequences early in training and getting more experience with the task. Third, Gi-DREADD activation did not alter goal-directed control of the previously reinforced sequence, as measured by the sequence rate and the latency to initiate left-leading sequences during selective satiation tests. Fourth, and finally, Gi-DREADD activation did not disrupt the insensitivity of completion times to outcome devaluation. Interpretation of these results are deferred to the *General Discussion*.

## 4. Experiment 3 (inhibiting D2 MSNs in the DMS)

### 4.1 Methods

#### 4.1.1 Subjects

Forty naïve Long-Evans rats (20 males and 20 females) were housed in identical conditions as rats in Experiments 1 and 2. Each rat was bred by crossing a D2 Cre transgenic male (source: Rat Resource & Research Center P40OD011062) with a wildtype female (source: Charles River Laboratories). Roughly half of all offspring were confirmed to express Cre in D2 dopamine receptor-expressing neurons (genotyping outsourced to Transnetyx). Both Cre positive (*n* = 20) and Cre negative (*n* = 20) rats were used in this experiment.

#### 4.1.2 Apparatus

The apparatus was identical to that used in Experiments 1 and 2.

#### 4.1.3 Surgery

Rats underwent stereotaxic surgery in which an AAV was bilaterally infused at the following coordinates (relative to bregma): AP, +0.7 mm; ML, +/-2 mm; DV, −5 mm (Paxinos & Watson, 2007). The method of surgery was the same as in Experiments 1 and 2. All rats received bilateral infusions of the AAV carrying the gene for the Gi-DREADD (AAV5-hSyn-DIO-hM4Di-mCherry), counterbalanced with sex and lineage.

#### 4.1.4 Behavioral training

Rats were trained on the same action sequence task as that used in Experiments 1 and 2 for 20 daily sessions, beginning a minimum of 3 weeks following surgery. There were four groups: DREADD+CNO (*n* = 10, 5 male and 5 female), DREADD+vehicle (*n* = 10, 5 male and 5 female), noDREADD+CNO (*n* = 10, 5 male and 5 female), and noDREADD+vehicle (*n* = 10, 5 male and 5 female). Pellet assignment, group assignment, and sex were counterbalanced.

#### 4.2.5 Behavioral testing

Expression tests and devaluation tests proceeded exactly as in Experiments 1 and 2.

#### 4.2.6 Histology

Rats were perfused and brains were sectioned and imaged exactly as in Experiments 1 and 2.

### 4.3 Results

#### 4.3.1 Histology

We observed robust mCherry expression in cell bodies within the DMS (Figure 6A, left). We also observed anterograde mCherry expression in axon terminals within the globus pallidus (Figure 6A, middle). Two rats, both in the DREADD+CNO group, were excluded from all analyses because fluorescence extended into the DLS. The boundaries of striatal fluorescent expression for all other rats are presented in Figure 6A.

**Figure 6.**
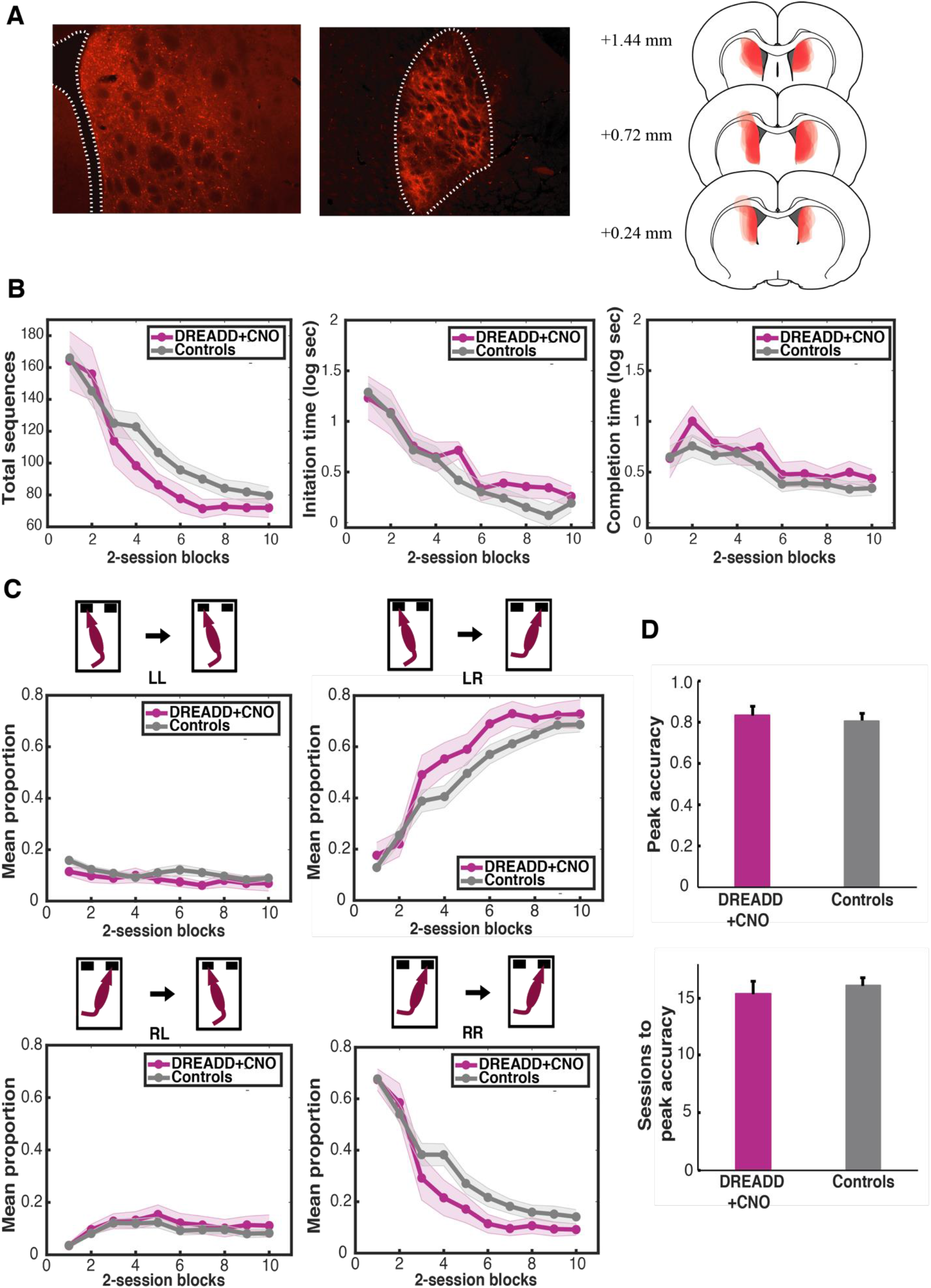
Training data from Experiment 3 (D2 DMS). (**A**) Left: Example coronal section showing mCherry expression in the DMS. The lateral ventricle is outlined. Middle: Coronal section showing axon terminal mCherry expression in the globus pallidus. Right: mCherry expression boundaries across all rats given DREADD AAV infusions in the DMS. (**B**) Total sequences (left), initiation times (middle), and completion times (right) across 2-session blocks for rats expressing DREADDs in the DMS and injected with CNO every day prior to training, and controls rats, Latency measures are averaged across all sequence types performed within a session. (**C**) Proportions of each sequence type across 2-session blocks. (**D**) Top: Mean peak accuracies for DREADD+CNO and control rats, defined as the maximum proportion of LR sequences achieved in a single session. Bottom: Mean sessions to peak accuracy.

#### 4.3.2 Training and expression testing

We examined measures of performance during training by analyzing the total number of sequences performed during each session, and the latency to initiate and complete all sequences during each session (Figure 6B). The data were once again collapsed across the three control groups, as there were no statistically significant differences detected among the means (between-group ANOVAs performed on every 2-session block: *F*’s < 1.17, *p*’s > .05). There were no differences detected between DREADD+CNO and control rats during any 2-session block for total sequences performed (MSE = 1392.60, *F*’s(1,22) < 2.71, *p*’s > .05), initiation times (MSE = 0.32, *F*’s(1,74) < 1.70, *p*’s > .05), or completion times (MSE = 0.22, *F*’s(1,100) < 0.38, *p*’s > .05). These data indicate that, overall, Gi-DREADD activation did not affect behavioral performance at any point during training.

To assess sequence acquisition, the relative proportions of each sequence type were examined (Figure 6C). Data from the three control groups were collapsed, as there were no statistically significant differences detected among the mean LR proportions (between-group ANOVAs performed on every 2-session block: *F*’s(2,65) < 2.20, *p*’s > .05). When comparing group means at each training block, significant differences were detected during block 4 for LR and RR sequences (LR: MSE = 0.04, *F*(1,96) = 3.98, *p* < .05; RR: MSE = 0.04, *F*(1,105) = 5.62, *p* < .05), with the DREADD+CNO group showing greater accuracy. No significant differences were detected with respect to LL or RL sequences (LL: MSE = 0.01, *F*’s(1,136) < 2.62, *p*’s > .05; RL: MSE = 0.01, *F*’s(1,129) < 0.77, *p*’s > .05). Groups did not differ with respect to peak accuracy (*t*(36) = 0.42, *p* > .05) or number of sessions to reach peak accuracy (*t*(36) = 0.49, *p* > .05; Figure 6D). These analyses suggest that Gi-DREADD activation in the DMS facilitated, by some measures, the learning of a reinforced action sequence.

During tests of expression conducted after training, there were no detectable within- or between-group differences between CNO and vehicle tests for either LR or RR proportions (*F*’s < 2.43, *p*’s > .05). We conclude that D2 MSNs in the DMS do not contribute to the expression of action sequence learning.

#### 4.3.3 Devaluation tests

Sensitivity to devaluation was first assessed by examining the target sequence rate under the four testing conditions: CNO/valued, CNO/devalued, vehicle/valued, and vehicle/devalued (Figure 7A). As in Experiments 1 and 2, the animals generally displayed a devaluation effect that, itself, was unaffected by the various training and test conditions. There were no significant differences between groups during any of the tests (MSE = 16.80, *F*’s(3,107) < 1.13, *p*’s > .05). Collapsing across groups, there was a significant effect of test (MSE = 11.73, *F*(3,102) = 20.77, Δ = 58.09, *p* < .05). Post-hoc contrasts revealed that rats performed fewer sequences during devalued than valued test sessions after CNO injections (*F*(3,102) = 10.81, *p* < .05) and vehicle injections (*F*(3,102) = 8.93, *p* < .05). Thus, Gi-DREADD activation in the DLS did not disrupt goal-directed control of sequences, as measured by the target sequence rate.

**Figure 7.**
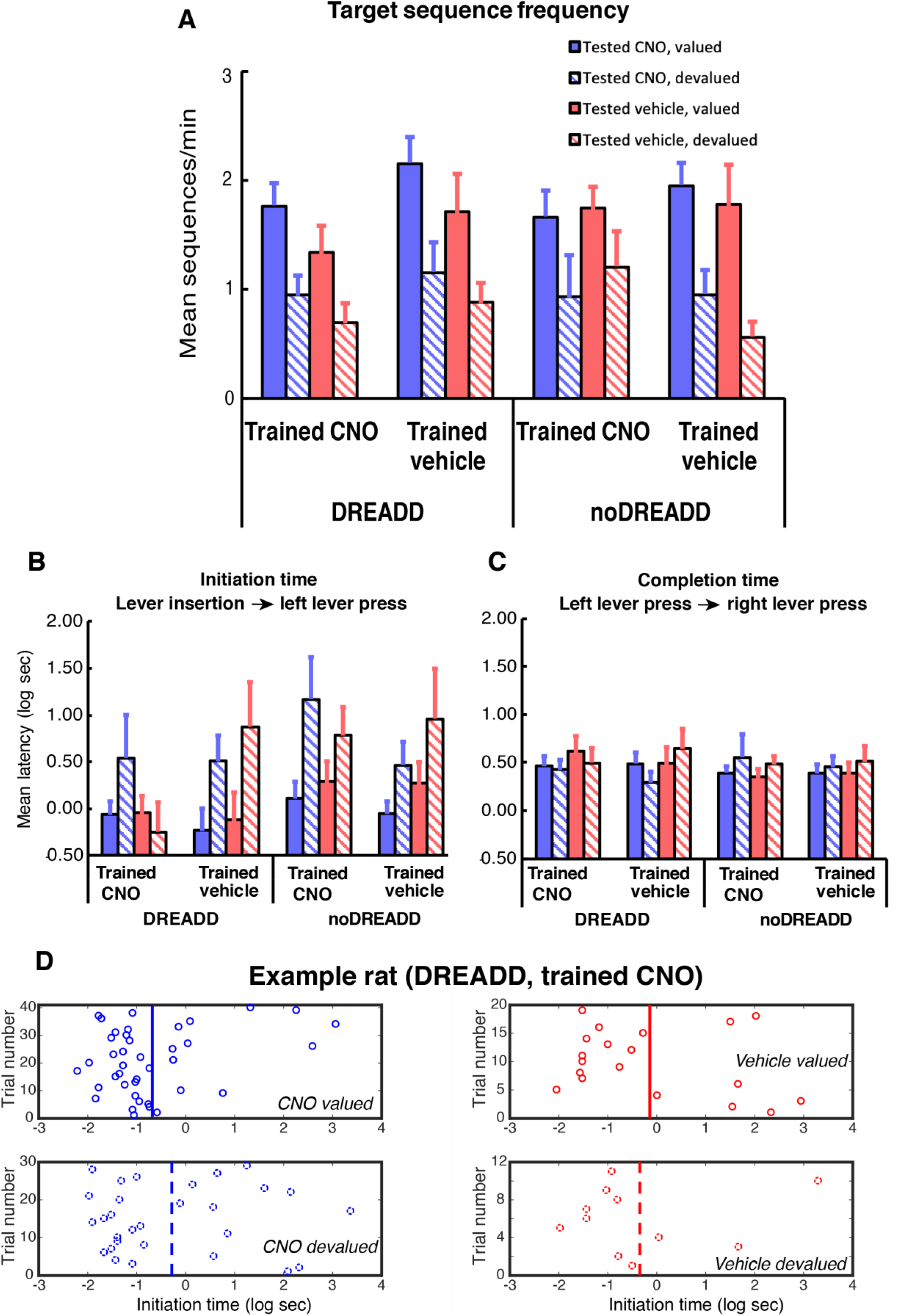
Devaluation test data from Experiment 3 (D2 DMS). (**A**) The rate of LR sequences during devaluation tests. (**B**) Initiation latencies during devaluation tests, defined as the time from lever insertion to the first left lever press. See panel A for legend. (**C**) Completion latencies during devaluation tests, defined as the time from a left lever press to a right lever press during LR trials. See panel A for legend. (**D**) An example rats from the DREADD+CNO training group, showing trial-by-trial left lever initiation times. Vertical lines represent session means.

We then analyzed latencies to initiate and complete sequences during devaluation tests as in Experiments 1 and 2 (Figure 7B). CNO and vehicle tests were once again analyzed separately. The four training groups displayed a similar pattern of data, i.e. faster initiation times during valued than devalued sessions, during CNO tests, but the devaluation effect was lost for the DREADD+CNO group when tested under vehicle conditions. A significant group difference was detected during the vehicle devalued tests (MSE = 1.09, *F*(3,55) = 2.23, Δ = 3.45, *p* < .05), and post-hoc contrasts confirmed that the DREADD+CNO training group showed faster initiation times compared to all other training groups (*F*(1,31) = 2.17, *p* < .05). A representative set of individual rat data is shown in Figure 12D. No between-group differences were detected during the three other tests (*F*’s < 1.37, *p*’s > .05). Furthermore, collapsing across groups, there were significant differences between valued and devalued test sessions (i.e. there was a main effect of test session), with rats being slower, overall, to initiate sequences during devalued than valued test sessions (CNO: MSE = 0.50, *F*(1,32) = 19.47, Δ = 17.25, *p* < .05; Vehicle: MSE = 0.68, *F*(1,31) = 7.47, Δ = 5.99, *p* < .05). This pattern of statistical results implies (Rodger, 1974) that initiation times were slower during devalued test sessions compared to valued test sessions following CNO injections for all groups (effect size = 0.80*σ*), and during vehicle test sessions for DREADD+vehicle, noDREADD+CNO, and noDREADD+vehicle training groups (effect size = 0.66*σ*). In contrast, the DREADD+CNO training group showed a slightly reversed effect with quicker mean initiation times during devalued than valued tests following vehicle injections (effect size = 0.13*σ*). These analyses show that removal of Gi-DREADD-mediated inhibition of D2 neurons in the DMS (following training with such inhibition) resulted in a disruption of goal-directed sequence initiation.

For completion latencies (Figure 7C), no between-group differences were detected during any of the tests (CNO: MSE = 0.15, *F*’s(3,52) < 0.69, *p*’s > .05; Vehicle: MSE = 0.19, *F*’s(3,47) < 0.62, *p*’s > .05), and, collapsing across groups, there were no significant differences between valued and devalued test sessions (CNO: MSE = 0.08, *F*(1,31) = 0.00, *p* > .05; Vehicle: MSE = 0.08, *F*(1,31) = 1.52, *p* > .05). These analyses show that chemogenetic inhibition spared the insensitivity of sequence completion times to outcome devaluation.

Consumption data from the satiation periods showed that all groups consumed the same amount of pellets across the four different test days (MSE = 1.13, *F*’s(3,102) < 1.50, *p*’s > .05). There were no between-group differences in overall intake collapsed across tests (MSE = 33.41, *F*(3,34) = 0.20, *p* > .05). Preference scores were analyzed as in Experiments 1 and 2. Within-group ANOVAs revealed no differences between preference scores on CNO and vehicle tests (MSE = 0.01, *F*’s(1,34) < 0.52, *p*’s > .05). However, a significant between-group difference when collapsing across CNO and vehicle tests was observed (MSE = 0.01, *F*(3,34) = 5.77, Δ =13.30, *p* < .05), and post-hoc contrasts showed that the DREADD+vehicle training group showed an overall lower preference score compared to all other groups. Collapsing across CNO and vehicle tests, the mean preference scores for the DREADD+CNO, DREADD+vehicle, mCherry+CNO, and mCherry+vehicle groups were 99%, 89%, 95%, and 96%, respectively.

### 4.4 Discussion

We sought to determine whether D2 MSNs in the DMS are necessary for action sequence learning and performance. There were five main findings. First, Gi-DREADD activation facilitated the rate at which a reinforced sequence was acquired, and this was true despite the absence of any evidence showing that Gi-DREADD activation sped up sequence performance. Second, Gi-DREADD activation did not appear to disrupt the goal-directed control of the previously reinforced action sequence in terms of the rate at which it was performed during reward devaluation. Third, Gi-DREADD activation during training altered learning in such a way that, when DREADD activation was removed during devaluation tests, the normal devaluation effect on sequence initiation times was lost. Fourth, Gi-DREADD activation did not disrupt the insensitivity of completion times to outcome devaluation. Interpretation of these results are deferred to the *General Discussion*.

## 5. Experiment 4 (inhibiting D2 MSNs in the DLS)

### 5.1 Methods

#### 5.1.1 Subjects

Thirty-nine naïve Long-Evans rats (25 males and 14 females) were housed in identical conditions as rats in Experiments 1-3. Each rat was bred by crossing a D2 Cre transgenic male (source: Rat Resource & Research Center P40OD011062) with a wildtype female (source: Charles River Laboratories). Roughly half of all offspring were confirmed to express Cre in D2 dopamine receptor-expressing neurons (genotyping outsourced to Transnetyx). Both Cre positive (*n* = 20) and Cre negative (*n* = 19) rats were used in this experiment.

#### 5.1.2 Apparatus

The apparatus was identical to that used in Experiments 1-3.

#### 5.1.3 Surgery

Rats underwent stereotaxic surgery in which an AAV was bilaterally infused at the following coordinates (relative to bregma): AP, +0.7 mm; ML, +/-3.6 mm; DV, −5 mm (Paxinos & Watson, 2007). The method of surgery was the same as in Experiments 1-3. All rats received bilateral infusions of the AAV carrying the gene for the Gi-DREADD (AAV5-hSyn-DIO-hM4Di-mCherry), counterbalanced with sex and lineage.

#### 5.1.4 Behavioral training

Rats were trained on the same action sequence task as that used in Experiments 1-3 for 20 daily sessions, beginning a minimum of 3 weeks following surgery. There were four groups: DREADD+CNO (*n* = 10, 6 male and 4 female), DREADD+vehicle (*n* = 10, 7 male and 3 female), noDREADD+CNO (*n* = 10, 6 male and 4 female), and noDREADD+vehicle (*n* = 9, 6 male and 3 female). Pellet assignment, group assignment, and sex were counterbalanced.

#### 5.1.5 Behavioral testing

Expression tests and devaluation tests proceeded exactly as in Experiments 1-3.

#### 5.1.6 Histology

Rats were perfused and brains were sectioned and imaged exactly as in Experiments 1-3.

### 5.2 Results

#### 5.2.1 Histology

We observed robust mCherry expression in cell bodies within the DLS (Figure 8A). Two rats, both in the DREADD+CNO group, were excluded from all analyses because fluorescence extended into the DMS. The boundaries of striatal fluorescent expression for all other rats are presented in Figure 7A.

**Figure 8.**
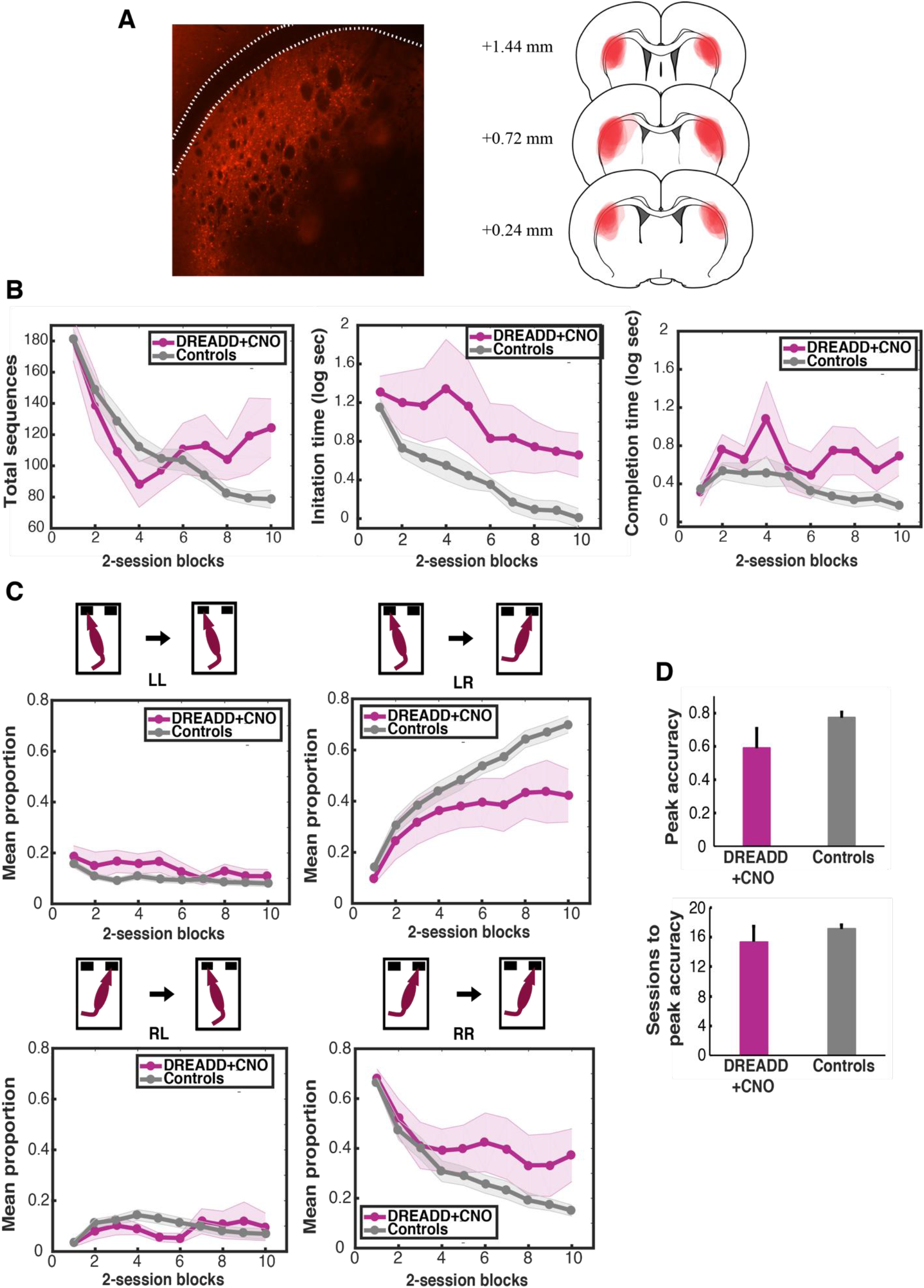
Training data from Experiment 4 (D2 DLS). (**A**) Left: Example coronal section showing mCherry expression in the DMS. The lateral ventricle is outlined. Right: mCherry expression boundaries across all rats given DREADD AAV infusions in the DLS. (**B**) Total sequences (left), initiation times (middle), and completion times (right) across 2-session blocks for rats expressing DREADDs in the DLS and injected with CNO every day prior to training, and controls rats. Latency measures are averaged across all sequence types performed within a session. (**C**) Proportions of each sequence type across 2-session blocks. (**D**) Top: Mean peak accuracies for DREADD+CNO and control rats, defined as the maximum proportion of LR sequences achieved in a single session. Bottom: Mean sessions to peak accuracy.

#### 5.2.2 Training and expression testing

We examined measures of performance during training by analyzing the total number of sequences performed during each session, and the latency to initiate and complete all sequences during each session (Figure 8B). The data were once again collapsed across the three control groups, as there were no statistically significant differences detected among the means (between-group ANOVAs performed on every 2-session block: *F*’s < 2.22, *p*’s > .05). Compared to control rats, the DREADD+CNO group performed significantly more sequences during the final two blocks of training (MSE = 1473.31, *F*’s(1,213) > 6.74, Δ’s > 5.68, *p*’s < .05). The latency to initiate sequences was also greater for the DREADD+CNO group compared to controls during blocks 3-5 and 7-10 (Figure 13B; MSE = 0.46, *F*’s(1,72) > 4.00, Δ’s > 2.89, *p*’s < .05), as were sequence completion times during blocks 4, 7, 8, and 10 (Figure 13B; MSE = 0.28, *F*’s(1,102) > 5.23, Δ’s > 4.13, *p*’s < .05). These data indicate that, overall, Gi-DREADD activation slowed sequence performance over the course of training, while also increasing the total number of sequences produced by the end of training.

To assess sequence acquisition, the relative proportions of each sequence type were examined (Figure 8C). Data from the three control groups were collapsed, as there were no statistically significant differences detected among the mean LR proportions (between-group ANOVAs performed on every 2-session block: *F*’s(2,55) < 2.27, *p*’s > .05). When comparing group means at each training block, significant differences were detected during blocks 3 and 5 for LL sequences (MSE = 0.01, *F*’s(1,120) > 5.61, Δ’s > 4.52, *p*’s < .05), blocks 7-10 for LR sequences (MSE = 0.04, *F*’s(1,69) > 5.51, Δ’s > 4.35, *p*’s < .05), and blocks 6, 7, and 10 for RR sequences (MSE = 0.04, *F*’s(1,71) > 4.00, Δ’s > 2.89, *p*’s < .05), with the DREADD+CNO group showing relatively poor task accuracy. No significant differences were detected with respect to RL sequences (MSE = 0.01, *F*’s(1,137) < 3.22, *p*’s > .05). Groups differed with respect to peak accuracy (*t*(35) = 2.16, *p* < .05) but not number of sessions to reach peak accuracy (*t*(35) = 1.14, *p* > .05; Figure 8D). These data indicate that Gi-DREADD activation during training resulted in a thwarted ability to acquire a sequence of lever presses.

One pressing question is whether the learning deficit that resulted from DREADD activation was a consequence of the DREADD activation negatively impacting the sequence learning process or was a by-product of a performance deficit. DREADD activation also resulted in a slowing down of sequence initiation and completion latencies across all sequences (Figure 8B), and this could have potentially reduced the number of learning opportunities throughout training. Alternatively, latencies to initiate and complete sequences could have stemmed from an underlying learning deficit, wherein poor learning causes a loss in motivation and a slowing down of behavior generally. Both of these possibilities seem likely, and the data cannot distinguish between either account. While the greater total number of sequences performed in the DREADD+CNO group at the end of training (Figure 8B) could be used to argue against a general performance deficit, that effect most likely originates from a combination of DREADD+CNO rats taking longer to achieve the 50 reward limit and also performing with lower accuracy.

During tests of expression that were conducted after training, there were no detectable within-group differences between CNO and vehicle tests for LR proportions (MSE = 0.01, *F*’s(1,33) < 0.67, *p*’s > .05). A between-group ANOVA revealed overall group differences (MSE = 0.04, *F*(3,33) = 2.47, *p* < .05), with the DREADD+CNO group performing with lower accuracy than the DREADD+vehicle and noDREADD+CNO groups, which collectively performed with lower accuracy than the noDREADD+vehicle group. We conclude that DREADD activation during training interfered with action sequencing, but this was not an issue of expressing latent learning.

#### 5.2.4 Devaluation tests

As was observed in Experiments 1-3 all training groups were equally sensitive to reward devaluation in vehicle and CNO test sessions (Figure 9A), with more LR sequences produced during valued than devalued tests. There were no differences between groups during any of the tests (MSE = 30.52, *F*’s(3,105) < 1.36, *p*’s > .05), but collapsing across groups, there were significant differences across the four test conditions (MSE = 21.47, *F*(3,99) = 7.54, Δ = 19.16, *p* < .05). Post-hoc contrasts revealed that rats performed fewer sequences during devalued than non-devalued test sessions after CNO injections (*F*(3,99) = 4.00, *p* < .05) and after vehicle injections (*F*(3,99) = 3.38, *p* < .05). Thus, although Gi-DREADD activation in the DLS impaired overall target sequence execution and/or learning, it did not disrupt goal-directed control of those sequences, as measured by the target sequence rate.

**Figure 9.**
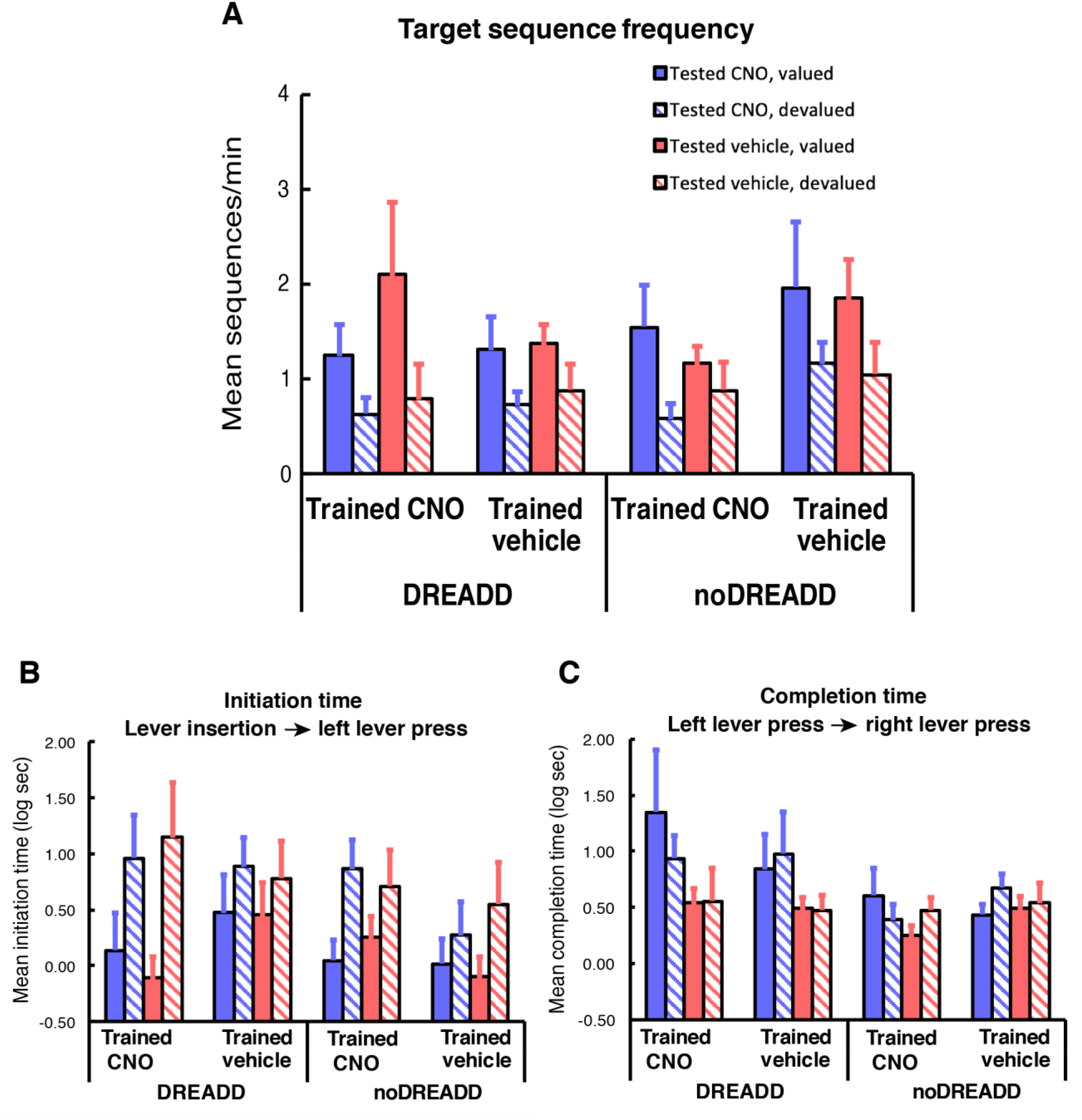
(**A**) The rate of LR sequences during devaluation tests in Experiment 4 (D2 DLS). (**B**) Initiation latencies during devaluation tests, defined as the time from lever insertion to the first left lever press. See panel A for legend. (**C**) Completion latencies during devaluation tests, defined as the time from a left lever press to a right lever press during LR trials. See panel A for legend. (**D**) An example rats from the DREADD+CNO training group, showing trial-by-trial left lever initiation times.

We then analyzed latencies to initiate and complete sequences during devaluation tests (Figure 9B). Unlike Experiments 1-3, CNO and vehicle tests were not analyzed separately because all rats performed at least one left-leading sequence during all test sessions. Generally speaking, all groups displayed longer initiation latencies on devalued than valued tests and this was equally true for CNO and Vehicle test sessions. There were no differences between groups during any of the tests (MSE = 0.82, *F*’s(3,96) < 1.10, *p*’s > .05), but collapsing across groups, there were significant differences amongst the four test sessions (MSE = 0.53, *F*(3,99) = 8.62, Δ = 22.34, *p* < .05). Post-hoc contrasts revealed that rats were slower to initiate left-leading sequences during devalued than valued test sessions after CNO injections (*F*(3,99) = 3.84, *p* < .05) and after vehicle injections (*F*(3,99) = 4.78, *p* < .05). This analysis further confirms that the chemogenetic manipulation did not affect goal-directed control of sequences, as measured by initiation latency.

For completion latencies (Figure 9C), CNO and vehicle tests were analyzed separately because some rats did not perform an LR sequence during one of the test sessions. There were no between-group differences detected during any type of test session (CNO: MSE = 0.75, *F*’s(3,57) < 1.73, *p*’s > .05; Vehicle: MSE = 0.19, *F*’s(3,64) < 0.79, *p*’s > .05), and, collapsing across groups, there were no significant differences between valued and devalued test sessions (CNO: MSE = 0.55, *F*(1,30) = 0.05, *p* > .05; Vehicle: MSE = 0.20, *F*(1,32) = 0.40, *p* > .05). These analyses show that chemogenetic inhibition spared the insensitivity of sequence completion times to outcome devaluation.

Consumption data from the satiation periods showed that all groups consumed the same amount of pellets across the four different test days (MSE = 13.82, *F*’s(3,99) < 0.95, *p*’s > .05), and there were no between-group differences in overall intakes collapsed across test days (MSE = 24.79, *F*(3,32) = 0.61, *p* > .05). Within-group ANOVAs on the preference score data revealed no differences between preference scores on CNO and vehicle tests (MSE = 0.01, *F*’s(1,33) < 3.71, *p*’s > .05), and there were no between-group differences either (MSE = 0.01, *F*(3,3) = 0.66, *p* > .05). Collapsing across CNO and vehicle tests, the mean preference scores for the DREADD+CNO, DREADD+vehicle, mCherry+CNO, and mCherry+vehicle groups were 96%, 99%, 97%, and 99%, respectively.

### 5.3 Discussion

We sought to determine whether D2 MSNs in the DLS are necessary for action sequence learning and performance. There were four main findings. First, Gi-DREADD activation slowed sequence performance over the course of training, as measured by the latency to initiate and complete sequences. Second, Gi-DREADD activation slowed the rate at which sequences were learned, but it is unclear whether this stemmed from a performance deficit or a learning deficit. Third Gi-DREADD activation did not alter goal-directed control of the previously reinforced sequence, as measured by the sequence rate and the latency to initiate sequences during selective satiation tests. This implies a distinction between learning a sequence and learning to respond in a goal-directed way. Fourth, Gi-DREADD activation did not disrupt the insensitivity of completion times to outcome devaluation.

## 6. General Discussion

Previous research implicates the dorsal striatum as playing a critical role in action sequence organization (see Garr, 2019 for review). The goal of this paper was to follow up on this work by chemogenetically inactivating neuronal subtypes in different regions of the dorsal striatum during and/or after action sequence acquisition. To achieve this goal, a virus carrying the gene for the hM4Di DREADD was infused locally into the either the DMS or DLS prior to behavioral training. The gene was subject to Cre-lox recombination such that the gene could only be transcribed in neurons containing Cre recombinase. Combining this technique with transgenic rats containing Cre only in either D1 or D2 dopamine receptor-expressing cells, the chemogenetic manipulations could be targeted specifically to dorsal striatal neurons that participate in the direct and indirect basal ganglia pathways.

Gi-DREADD function was validated by combining unilateral DREADD expression with systemic CNO and caffeine injections followed by c-Fos immunohistochemistry. Previous efforts to validate Gi-DREADD function in the striatum have relied mostly on slice electrophysiology, and have shown that CNO-mediated activation of DREADD-expressing neurons induces hyperpolarization and reduces the spike rate evoked by depolarizing currents (Dobbs et al., 2016; Ferguson et al., 2011; Francis et al., 2015; Zhang et al., 2018). Here, we used c-Fos expression as a proxy for neural activation, which is an approach that has previously been used with DREADDs in the dorsal striatum (Ferguson et al., 2011; Ferguson et al., 2013) and in brain areas outside the striatum (Kane et al., 2017; Keenan et al., 2017; Siegel et al., 2015). While the c-Fos results are consistent with CNO-mediated inhibition of neurons expressing Gi-DREADDs, the mechanism by which this inhibition works is unclear. The common explanation for how activation of the hM4Di DREADD inhibits neuronal firing is via activation of G-protein inwardly rectifying potassium channels (GIRKs; Roth, 2016). However, *in situ* hybridization studies show no GIRK expression in striatal neurons (Lein et al., 2007), and the mechanism by which Gi-DREADDs work in the striatum is currently a topic of ongoing investigation (Voyvodic, Abrahao, & Lovinger, 2018).

Previous research with mice has provided evidence for different contributions of striatal subregions and neuronal subtypes in action sequence acquisition. In one experiment, pre-training excitotoxic lesions of the DLS slowed down the rate of learning a left-right lever press sequence (Yin, 2010). Mice given DMS lesions, however, did not show a learning impairment and behaved liked sham controls. Another experiment that employed the same task focused exclusively on the DLS but used a permanent inactivation method to inhibit MSNs participating in the direct and indirect pathways by using D1 and A2A Cre mice, respectively (Rothwell et al., 2015). Inactivation of direct pathway, but not indirect pathway, MSNs disrupted the acquisition of the reinforced sequence, and the same impairment in task accuracy was also observed when inactivation was introduced after task acquisition. We were unable to replicate this finding, and instead showed that inactivating indirect pathway MSNs in the DLS impaired sequence acquisition (see Experiment 4).

What accounts for this critical difference between studies? While we used temporary chemogenetic inactivations, Rothwell and colleagues (2015) virally expressed inwardly rectifying potassium channels, which create a chronically high action potential firing threshold (Lin et al., 2010; Rothwell et al., 2014). While this fact makes it difficult to compare our studies, a key piece of missing information from the study by Rothwell and colleagues (2015) is latency data. Perturbation of striatal neurons forming the direct pathway has often been shown to slow down movement execution while perturbation of neurons forming the indirect pathway often results in hyperactivity (e.g. Drago et al., 1998; Durieux et al., 2009; Kravitz et al., 2010; Panigrahi et al., 2015; Sano et al., 2003). This is often described as the Go/No-Go model of the direct and indirect pathways (Bariselli et al., 2018). Although our data did not generally conform to this model, if A2A Cre mice were indeed hyperactive with shortened response latencies, this could have sped up sequence learning when it otherwise could have been impaired. This is because shorter action latencies could have allowed for more learning opportunities. Similarly, the slowing down of sequence acquisition observed in D1 Cre mice could have stemmed from a slowing down of action latencies. Because latency data were not reported by Rothwell et al. (2015), it is difficult to know whether performance speed or the different neural manipulations employed in our studies explains our different results.

As noted above, rather than observing a clear retardation and facilitation of performance speed when inhibiting D1 and D2 MSNs, respectively, we observed a pattern of results that was not fully consistent with the Go/No-Go model of dorsal striatal function. When inhibiting D1 MSNs in the DMS, we indeed observed a slowing down of task performance, both in the latency to complete sequences and the total number of sequences executed. This slowing down of performance was only observed early in training, consistent with prior accounts of the DMS contributing early but not late during skill learning (Ashby, Turner, & Horvitz, 2010; Miyachi et al., 1997; Miyachi, Hikosaka, & Lu, 2002; Yin et al., 2009). Performance speed was not affected when inhibiting D2 MSNs in the DMS. However, the data from the DLS studies directly contradict the Go/No-Go model of the direct and indirect pathway function. When inhibiting D1 MSNs in the DLS, we observed a facilitation of sequence performance early in training, while inhibiting D2 MSNs in the DLS resulted in a slowing of performance that was likely due to impaired learning. This disparity is not surprising given the number of studies in recent years that have reported inconsistencies between theory and data. For example, one study showed that Gi-DREADD activation in the DLS of young D1 Cre mice resulted in faster rates of lever pressing during a free operant random ratio task (Matamales et al., 2017). Another study showed that optogenetically inhibiting D2 MSNs in the DLS during fixed ratio lever pressing slowed down the rate of pressing, as did high frequency stimulation of D1 MSNs (Tecuapetla et al., 2016). Notably, these experiments, like the ones reported here, were conducted in the context of motivated reward seeking, while many of the studies that support the Go/No-Go model are studies of spontaneous movement in open arenas (Bateup et al., 2010; Durieux et al., 2009; Kravitz et al., 2010; Ryan et al., 2018).

The finding that inhibition of D2 MSNs in the DLS interfered with the acquisition of an action sequence is consistent with other studies of motor learning. For example, in one study mice were trained to press levers in a left-left-right-right pattern while optogenetic stimulation was applied to either D1 or D2 MSNs (Geddes et al., 2018). Stimulating D2 MSNs immediately prior to the execution of the first left lever press in the sequence caused the animal to immediately switch to the right lever before even executing the left lever press, as did stimulating during the execution of the first press. Electrophysiological recordings also revealed that a high proportion of D2 MSNs increased their firing rates between the left and right subsequences, suggesting that these neurons contribute to switching between actions in a sequence. Consistently, inhibiting D2 MSNs in the DLS (Experiment 4) led to a reduction in LR sequences, as well as an increased probability of performing LL and RR sequences. In another motor learning study using the accelerated rotarod in mice, patch-clamp recordings revealed that D2 MSNs in the DLS, but not D1 MSNs, underwent significant potentiation of excitatory transmission following eight days of training compared to naïve mice (Yin et al., 2009). These findings collectively suggest that the indirect pathway originated from the DLS is an important circuit for skilled motor learning, and may specifically contribute to switching between actions within a complex sequence.

Following action sequence acquisition, we conducted tests of goal-directed control by sating rats either on the pellet type associated with LR sequence execution or another control pellet type of a different flavor. Across all experiments, goal-directed control of the rate of the reinforced action sequence was immune to chemogenetic inhibition. This is especially surprising for the experiments in which DREADDs were expressed specifically in the DMS, given that prior research has consistently shown that lesions and blockade of synaptic plasticity in the DMS erase goal-directed control of lever pressing rates in free operant tasks (Gremel & Costa, 2013; Shiflett, Brown, & Balleine, 2010; Yin et al., 2005a, 2005b). However, the lever sequence task used in the present experiments differs in a potentially crucial way from the free operant, single lever tasks used in prior research on striatal control of goal-directed behavior. The task used in the present set of experiments continuously reinforced a single sequence, while in most studies of goal-directed control it is common to use partial reinforcement schedules in which the timing of the reward is uncertain and there are often many responses emitted between reward deliveries. In contrast, the sequence task employed in the present set of experiments was associated with a high degree of certainty regarding the timing of rewards, as well as relatively short latencies separating actions from rewards. It has been hypothesized that these variables—temporal certainty of outcomes and action-outcome contiguity—play important roles in determining the goal-directed nature of behavior (DeRusso et al., 2010; see also Garr et al., in press), and it is possible that tasks that maximize these variables, such as the sequence task used here, could prevent goal-directed control from ever being erased (however, see Adams, 1982). On the other hand, it has been emphasized that it is the posterior DMS specifically that participates in goal-directed decision-making (Peak, Hart, & Balleine, 2018), while the virus infusions here were aimed at the anterior DMS. However, there is some evidence that disruption of anterior DMS function does interfere with goal-directed control of instrumental actions (Corbit, Nie, & Janak, 2012).

Despite no disruptions of goal-directed control when measuring LR sequence rates, we did find that chemogenetic inhibition during training disrupted aspects of sequence initiation and completion during devaluation tests. Generally, and consistent with our previous results (Garr & Delamater, 2019), initiation latencies were sensitive to reward devaluation while completion latencies were not following a moderate amount of training. However, D1 Cre rats that previously experienced Gi-DREADD activation in the DMS during training slowed the time to complete LR sequences during reward devaluation, but only when DREADDs were active (Experiment 1). This result suggests that, under normal circumstances, the direct pathway via the DMS thwarts the development of goal-directed sequence completion during learning. Why a similar slowing of completion latencies was not observed during the vehicle tests is puzzling, but can be explained by the possibility that removal of Gi-DREADD activation disrupted the retrieval of prior learning. While this result is novel and perhaps unexpected, it is generally consistent with the identification of D1 MSNs as part of a ‘Go’ pathway, wherein the ‘Go’ signal is a signal to complete an action sequence regardless of the value of the consequent outcome. If the recruitment of D1 MSNs in the DMS during action sequence learning is necessary for the prevention of goal-directed sequence completion, then one prediction is that, during extended training when sequence completion transitions to becoming goal-directed (Garr & Delamater, 2019), these neurons should become downregulated. A consequence of this prediction is that stimulating these neurons after extended training should result in a return of completion latencies to devaluation insensitivity.

Another complex finding was that in D2 Cre rats that experienced Gi-DREADD activation in the DMS during training, initiation times became insensitive to reward devaluation when DREADD activation was removed (Experiment 4). This result suggests that when action sequence learning occurs in the presence of inactive D2 MSNs in the DMS, introducing D2 MSN activation after learning impairs the retrieval of goal-directed sequence initiation. A broader conclusion that can be drawn from this suggestion is that retrieving the knowledge required for goal-directed sequence initiation requires the relative quiescence of D2 MSNs in the DMS. If these neurons are overactive relative to the conditions under which sequence learning occurred, then retrieval will be impaired. However, if D2 MSNs are underactive or equally active relative to the conditions of sequence learning (as was true of the DREADD+vehicle training group in Experiment 4), then retrieval will not be impaired. One prediction is that stimulating the activity of these neurons after learning in the absence of stimulation should also impair goal-directed sequence initiation. This prediction could be addressed with the use of excitatory, rather than inhibitory, DREADDs. Another prediction is that D2 MSNs in the DMS should be recruited with extended training, since extensive training leads to abolished goal-directed control of sequence initiation (Garr & Delamater, 2019). A consequence of this prediction is that inhibiting these neurons after extended training should result in a return of sequence initiation to devaluation sensitivity.

In conclusion, transient inhibition of neuronal subtypes in the dorsal striatum largely spared the acquisition of an action sequence while having subtle effects on the sensitivity of sequence initiation and completion to outcome devaluation. Only chemogenetic inhibition of D2 receptor-expressing neurons in the DLS (but not D2 neurons in the DMS or D1 neurons in either the DMS or DLS) impaired sequence acquisition, and this finding supports the notion that the neural mechanisms that enable action sequence learning may reside specifically in the lateral portion of the dorsal striatum (Yin, 2010). In addition to largely sparing action sequence acquisition, the chemogenetic manipulations employed in the present set of studies completely spared goal-directed control of sequences as measured by the rate of sequence performance during outcome devaluation tests. This finding raises the question of whether the role of the DMS in goal-directed control of instrumental actions applies to tasks beyond free operant random ratio and random interval settings. Notably, a role for the DMS in goal-directed control of instrumental performance under free operant conditions has not yet been demonstrated using a non-specific chemogenetic approach, although it has been shown that DREADD-mediated inhibition of D2 MSNs in the DMS leaves goal-directed lever pressing unaffected in mice (Poyraz et al., 2016). Finally, we also show that while goal-directed control of sequence rates were immune to chemogenetic inhibition applied either during and/or after training, we found that learning to initiate and complete action sequences in a goal-directed manner depends, to some extent, on specific neural pathways originating from specific regions of the dorsal striatum. The direct and indirect pathways that originate in the DMS appear to play important roles in the goal-directed control of sequence completion and initiation, respectively. These findings highlight the importance of examining action sequences in terms of separate initiation and completion measures. Future work would benefit from using inhibitory DREADDs after more extensive training, as well as using alternate methods of circuit manipulation, such as optogenetics or even excitatory DREADDs. In addition, it would be beneficial to expand the study of action sequencing by adding more actions to the reinforced sequence. The left-right lever sequence task, while relatively easy to learn for rats, is limited in that sequence execution and completion are confounded. Lengthier sequences would provide an opportunity to study how manipulations of basal ganglia circuit function differentially affect actions in the middle of a sequence as opposed to actions more proximal to reward.

## Acknowledgements

This work was supported by a National Institute on Drug Abuse grant (SC1DA034995) awarded to ARD and a CUNY doctoral dissertation fellowship awarded to EG.

## Notes

#### Summary of Updates

Addition of missing reference (Geddes et al., 2018).

